# A single-cell and spatial atlas of prostate cancer reveals the combinatorial nature of gene modules underlying lineage plasticity and metastasis

**DOI:** 10.64898/2026.03.25.711335

**Authors:** Hanbing Song, Junxiang Xu, Keila Velazquez-Arcelay, Arda Demirci, Brendan L. Raizenne, Sarah C. Hsu, Joshua Choi, Julia H. Pham, Yih-An Chen, Hannah N. W. Weinstein, Isaac Salzman, Margaret Tsui, Jon Akutagawa, Wisdom Adingo, Ezequiel Goldschmidt, Peter R. Carroll, Julian C. Hong, Christopher M. Heaphy, Matthew R. Cooperberg, Nancy Greenland, Joshua D. Campbell, Franklin W. Huang

## Abstract

Prostate cancer encompasses a spectrum of disease states driven by complex cellular heterogeneity. To delineate the transcriptional programs underlying lineage plasticity and metastasis, we constructed a comprehensive single-cell atlas of 128 patients, spanning localized, castration-resistant, and metastatic disease. Lineage plasticity was prevalent in localized disease, with subsets of tumor cells adopting distinct basal-like and club-like states. Luminal-like cancer cells also displayed extensive lineage infidelity, defined not by a binary loss of identity but by the combinatorial erosion of luminal gene modules associated with higher grade and stage. In the metastatic setting, gene program association analysis (GPAS) identified a broad induction of cell-cycle gene modules across organ sites as well as an induction of organ-specific gene modules, including osteomimetic signaling in bone, neuro-migratory genes in brain, and erythroid-like transitions in liver. Neuroendocrine prostate cancers (NEPCs) were not monolithic but defined by combinations of NE-associated gene modules including a novel HES6 program. Notably, these modules were detected at intermediate levels in localized samples, suggesting molecular plasticity precedes histological transformation. We also developed a refined NE signature that could distinguish NEPC tumors more accurately than previously published signatures. Within the tumor microenvironment (TME), we observed an elevation of pro-inflammatory Th17 T-cells in African American patients and identified a rare Schwann cell population. Finally, we present PCformer, a transformer-based foundation model trained on >500,000 cells to automate cell-state classification. Together, this comprehensive atlas demonstrates the complex nature of gene modules underlying lineage infidelity and plasticity in cancer cells and highlights distinct immune and stromal populations within the tumor ecosystem.

## Introduction

Prostate cancer (PCa) is the second most diagnosed malignancy and a leading cause of cancer-related mortality in men globally, with over 1.4 million new cases annually^1^. While localized disease is often curable through surgery or radiation, metastatic castration-resistant prostate cancer (mCRPC) remains a lethal clinical challenge, largely due to its molecular heterogeneity, dynamic tumor microenvironment (TME), and complex treatment resistance mechanisms^2,3^. Recent studies leveraging single-cell and spatial transcriptomics^4–6^ have begun to reveal key cancer cell states or stromal and immune cell populations in the TME that contribute to disease progression and therapeutic resistance. The luminal epithelial cells are the putative cell of origin for prostate cancer and generally characterized by high expression of KLK3 (i.e. Prostate-specific antigen) with a dependency of the androgen receptor (AR) signaling. Distinct subtypes have been identified based on expression such as androgen receptor (AR)-high^7–10^, PSA-low/AR-low^11,12^, NE-like (neuroendocrine-like)^13^, double-negative (AR⁻/NE⁻)^14^ subtypes. In the setting of advanced disease, lineage plasticity to neuroendocrine phenotypes is associated with metastasis and resistance to androgen directed therapies^9,15^.

While bulk RNA sequencing methods have been primarily used to identify subtypes of disease resistant to therapy^16,17^, they are limited in their ability to fully capture the cellular diversity in the TME or transcriptional heterogeneity in cancer cells that drive cancer initiation, metastasis, and resistance to therapy. Single-cell RNA sequencing (scRNA-seq) has revolutionized our ability to resolve the cellular complexity of cancer tissue. Several studies have used scRNA-seq to reveal novel insights into PCa heterogeneity including characterizing immune infiltrates and stromal interactions that are associated with disease progression^18–20^. Despite these advances, a major limitation is that the overall number of metastatic samples is limited due to the challenges associated with obtaining biopsies from advanced-stage patients and metastatic sites. Furthermore, the number of metastatic tumor samples per organ site is even more limited further reducing statistical power to draw robust conclusions about organ-specific adaptations^18,21,22,23,24^. Beyond the total number of independent patients, the transcriptional heterogeneity that can be exist in the cancer cells within each sample adds additional complexity. Each cancer cell can have a unique combination of genetic alterations, epigenetic changes, and cell-cell interactions with the TME that influence its transcriptome and produce patient-specific clusters. New matrix factorization paradigms have been developed for detection of gene modules in heterogeneous cancer cells across tumors and studies and perform gene program association studies (GPAS) to determine how these modules are associated with important clinical phenotypes^25,26^. However, these approaches have yet to be applied to a large-scale atlas of prostate cancer cells.

To address the challenges of prostate cancer heterogeneity, we constructed a comprehensive single-cell atlas integrating data from 128 patients across localized and metastatic disease stages. By combining robust tumor identification, gene module analyses, we uncovered patterns of lineage plasticity and infidelity, including shared and organ-specific metastatic programs and ancestry-associated immune alterations. Furthermore, we developed PCformer, a transformer-based foundation model to facilitate reproducible cell-type annotation. Collectively, this atlas provides a unified framework for dissecting the molecular drivers of progression and lays the foundation for improved biomarker discovery and novel therapeutic development.

## Results

### Integration and tumor clustering of the PCa scRNA-seq atlas reveals basal-like and club-like tumor populations

A large-scale PCa single-cell RNA-seq atlas was developed by integrating 12 studies^18,20–22,24,27–33^ and 9 unpublished samples encompassing samples from 128 patients (**Supp Table 1)**. 95 samples were from patients with localized PC, 10 samples were from patients with CRPC (i.e. castration resistant but non-metastatic), and 23 samples were patients with metastatic CRPC (mCRPC) taken from 7 metastatic sites. After quality control and integration, 560,492 cells were annotated in 9 major cell populations, including epithelial, immune, stromal, and endothelial cells, with further sub-annotation of immune subsets, each showed enriched expression of canonical markers (**Figure 1a-b**, **Supp Figure 1a-c**). After sub-clustering of 263,017 epithelial cells, and luminal, basal, and club cells were identified using established signatures (**Supp Figure 1d**). 106,998 putative tumor cells were identified as those with substantial copy number changes above background (**Supp Figure 1e-g**). Integration and clustering of tumors cells was performed to reveal 15 broad clusters (**Figure 1c**). Different clusters showed enrichment of different cell type and pathway signatures (**Figure 1d**). Clusters 0, 1, 2, 5, and 12 contained higher levels of an AR signaling signature and were predominantly enriched in localized disease. Clusters 8 and 10 expressed a neuroendocrine signature and were predominantly found in metastatic samples from liver, lymph nodes, and CRPC prostate tissues. Cluster 14 was high for an epithelial-to-mesenchymal (EMT) signature defined by *VIM* and *GPM6B*, and was predominantly found in localized PCa samples. Some clusters were more strongly defined by biological processes distinct from canonical cell types. Clusters 6 and 10 were enriched for cell cycle pathways and represented cells undergoing mitosis while cluster 11 had higher levels of an inflammatory response signature defined by *CD69* and *CXCR4*.

**Figure 1.**
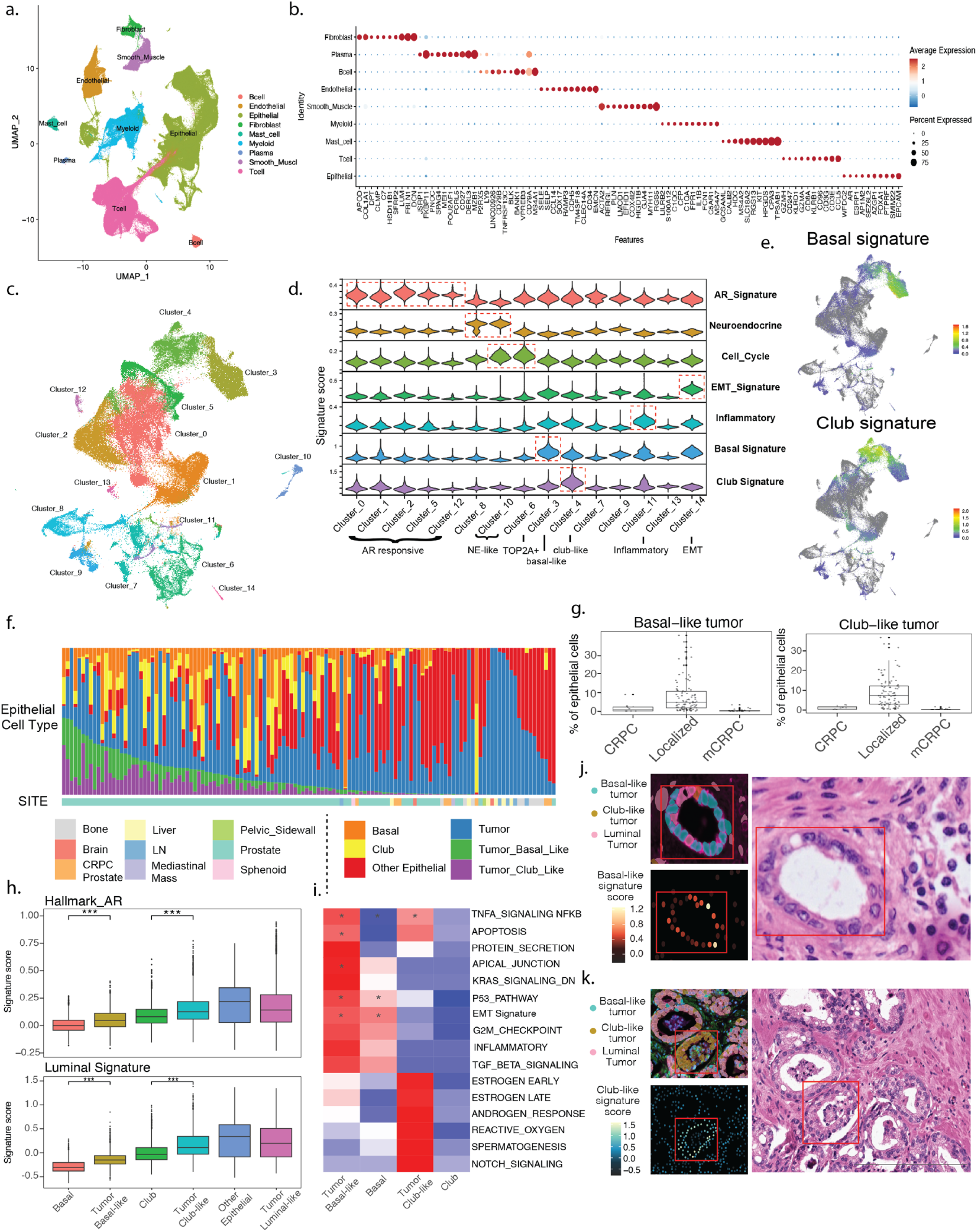
Tumor clustering of the PCa scRNA-seq atlas reveals tumor cell heterogeneity including basal-like and club-like tumor populations. **a.** Top-tier annotation of major cell types for the integrated PCa scRNA-seq atlas (N = 560,492 from 14 datasets). Cell type labels are color coded. **b.** Dot plot showing expression of canonical marker genes used to annotate major cell populations in the atlas. Dot size indicates the percentage of cells expressing each gene, and color intensity represents average expression. **c.** UMAP visualization of putative tumor cells (N = 106,998) identified based on elevated copy number alterations, showing clustering into 15 tumor cell clusters. Cluster identities are labeled. **d.** Stacked violin plots showing distribution of multiple prostate cancer–related gene signature scores across tumor clusters. **e.** Feature plots showing basal-like and club-like tumor signatures projected onto the tumor UMAP. **f.** Stacked bar plot showing the proportion of epithelial cell states across individual patients. Patients are ordered by the composition of basal and club-like tumor cells. **g.** Box plots showing the percentage of basal-like tumor (left) and club-like tumor (right) across localized PCa, CRPC prostate, and mCRPC samples. **h.** Box plots showing Hallmark androgen response and luminal signature scores across epithelial cell states. **i.** Heatmap showing pathway enrichment scores from gene set enrichment analysis (GSEA) comparing basal-like and club-like tumor cells with their benign counterparts, highlighting differentially enriched pathways. **j.** Spatial validation of basal-like tumor cells using one FFPE Human Prostate Adenocarcinoma with 5K Human Pan Tissue and Pathways Panel. Upper left image shows the spatial distribution of basal-like (cyan), club-like (gold), and luminal (pink) tumor cells within a representative region. Lower left image shows the basal-like tumor signature score for the same region. Right H&E image shows the corresponding histological features, with the red box highlighting a small region of a prostatic gland in a low-grade (Gleason 3) area where basal-like tumor signature is enriched. **k.** Spatial validation of basal-like and club-like tumor cells using a second FFPE Human Prostate Adenocarcinoma with 5K Human Pan Tissue and Pathways Panel. Upper left image shows the spatial co-localization of basal-like (cyan) and club-like (gold) tumor cells with other luminal tumor cells within a representative region. Lower left image shows the club-like tumor signature score for the same region. Right H&E image shows the corresponding histological features of the annotated region, with the red box highlighting tumor glands with amphophilic cytoplasm and prominent nucleoli where club-like signature is enriched.

Although luminal cells are the primary cell-of-origin for PCa, we observed that clusters 3 and 4 exhibited high levels of basal-like (*TP63*+) and club-like (*LTF*+, *LCN2*+) signatures, respectively (**Figure 1e**). Basal-like tumor cells represented 8.13% of all epithelial cells on average across localized PCa patients but only 1.87% in CRPC prostate tissues and 0.34% in mCRPC patients (p = 1.73e-11, Wilcoxon test; **Figure 1f,g**). Similarly, club-like tumors represented 8.86% of all epithelial cells on average across localized PCa patients but only 0.95% in CRPC prostate tissues and 0.15% in mCRPC patients (p = 2.2e-13, Wilcoxon test). These populations were predominantly found in localized PCa and were largely absent in advanced states, suggesting that the plasticity between basal and club-like states are progressively lost during disease progression. Compared to basal and club cells without detectable copy number alterations, the tumor basal and club-like cells showed higher levels of luminal and AR signatures (**Figure 1h**) but lower signatures compared to non-tumor luminal cells and the luminal-like tumor cells, suggesting that these basal-like and club-like tumor cells may represent specific tumor clusters with strong resemblance to the benign basal and club epithelial cells and simultaneously partial activation of androgen response pathway. Gene set enrichment analysis (GSEA) further revealed that basal-like tumor cells were enriched for pathways related to inflammation, and EMT signaling, while club-like tumor cells showed enrichment of androgen response, protein secretion, and oxidative stress pathways compared to their benign counterparts (**Figure 1i**), indicating that these two clusters represent distinct transcriptomic tumor profiles.

To understand the relationship between histological phenotypes and cancer cell states identified by scRNA-seq, we analyzed two localized PCa samples profiled with the Xenium 5K spatial transcriptomics platform (**Methods**). Tumor cell states were projected onto the spatial data using a label transfer method and each cell was scored using the established basal-like and club-like tumor state signature gene sets (**Methods**). In the first sample (**Supp Figure 1i**), several basal-like tumor cells were localized to a small region of a prostatic gland in a low-grade (Gleason 3) area (**Figure 1j**). The basal-like signature score was markedly enriched in this glandular structure, suggesting that cells in the basal-like tumor state are largely confined to well-differentiated glandular areas. In the second sample (**Supp Figure 1j**), both basal-like and club-like tumor cells were identified in tumor glands with amphophilic cytoplasm and prominent nucleoli (**Figure 1k**). Club-like tumor cells were the predominant population within the highlighted gland, and club-like signature scores were enriched. The spatial co-occurrence of these states within the same histological regions suggests that basal-like and club-like tumor states co-exist spatially and supports the presence of these tumor cell states identified in the scRNA-seq data.

### An atlas of gene modules in localized and metastatic PCa

As PCa has substantial intra- and inter-tumoral transcriptional heterogeneity, we sought to catalogue the combinations of gene modules that defined localized and metastatic samples. Celda^34^ was used to identify 166 co-expression modules (**Supp Table 2)** and linear mixed effect modules (LMEs) were used to subtract effects of different studies while maintaining biological variability across samples and disease sites (**Methods**; **Figure 2a**). Each module contained a group of genes with similar expression patterns across cells (i.e. co-expressed), while each module had a distinct pattern of expression across cells compared to other modules. The modules were annotated and divided into broad biological categories including those related to lineage, cell cycle and proliferation, extracellular matrix production (ECM)/epithelial-to-mesenchymal transition (EMT), antigen presentation, stress response, chemokine signaling, interferon signaling, and ribosomal/housekeeping. Examples of modules from different categories included the luminal modules L20:Luminal (*KLK2/3*) and L24:AR_Signaling (*AR*) (**Figure 2b**), an antigen presentation module was L104:MHC-I which contained major histocompatibility complex genes *HLA-A, HLA-B,* and *HLA-C*, a cell cycle module L156 enriched for cell cycle checkpoint genes, and immune-related modules enriched for interferon response genes such as L120:Interferon (*IFI6/35/44*) or signaling cytokines such as L100:Chemokine (*CXCL1/3*) (**Figure 2b**). When examining the percentage of variation explained by different factors, the modules most affected by variation between studies were the broader housekeeping modules enriched for ribosomal genes (Study Effect >40%; **Supp Figure 2a**). In contrast, modules affected by variation between metastatic sites were those related to other categories such as lineage and signaling (Site >20% **Supp Figure 2a**) suggesting that many modules are driven by biological variation despite the technical differences between studies.

**Figure 2.**
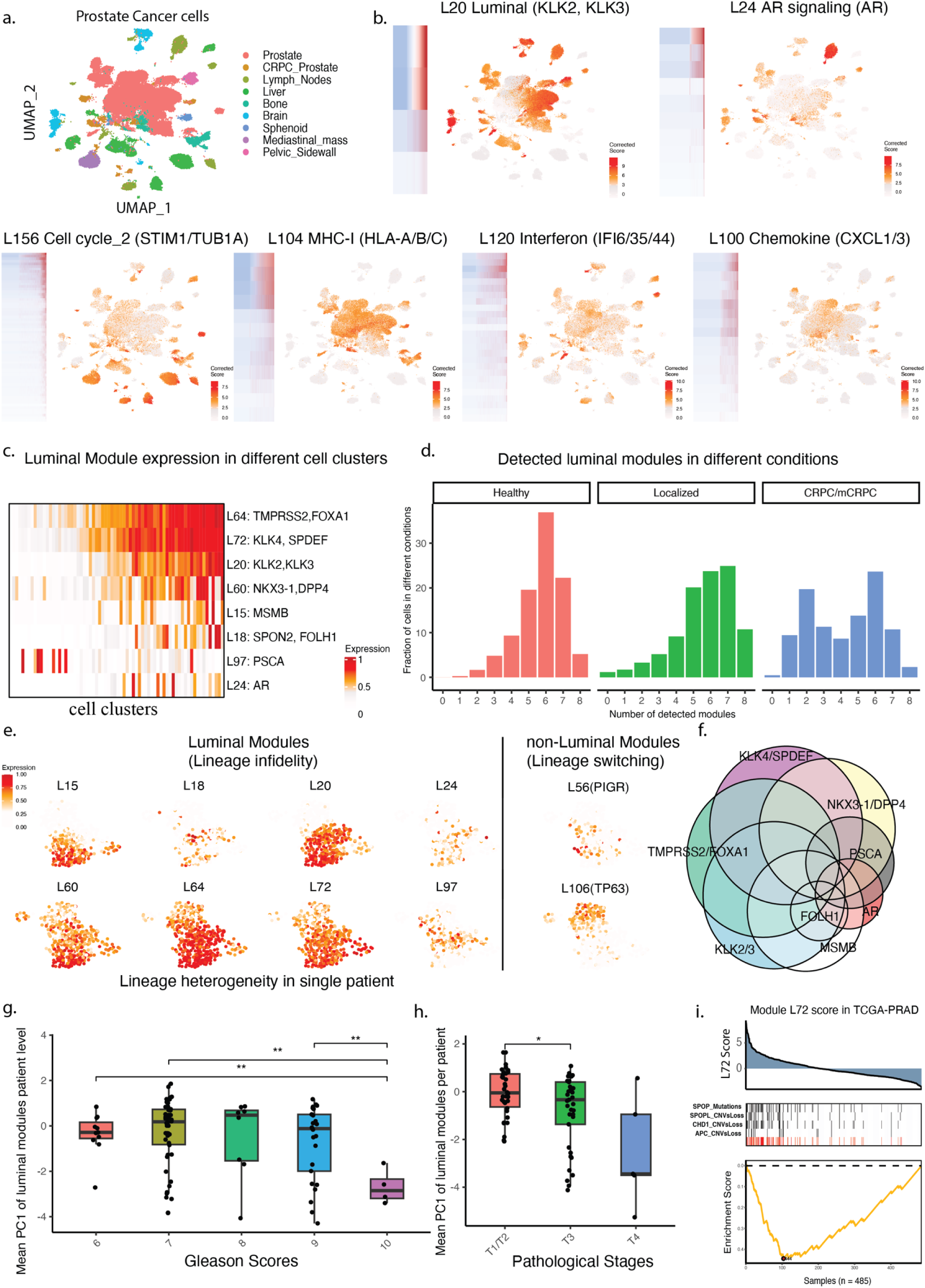
Gene module analysis highlights luminal lineage infidelity in prostate cancer. **a.** UMAP plot generated from module expression corrected for study-level effects but not tumor-level effects. Cells are colored by originating site. Cells from metastatic samples tend to form their unique own clusters. **b.** Examples of co-expression modules derived from Celda are shown. Each heatmap shows the co-expression pattern of genes assigned to that module. Cells in the heatmap are ordered by those with the lowest to the highest module probability. Each UMAP shows a unique pattern of log2 normalized module expression across cells. **c.** Heatmap shows the expression of modules containing previously described luminal markers across cell clusters. Each row is a module, and each column is a cell cluster from Celda. **d.** Each luminal module was classified as “detected” or not in each cell based on the module density. The percentage of cells in which a specific number of luminal modules were detected is shown for cells from healthy donors, localized PCa, and CRPC/mCRPC. Cancer cells from localized tumors exhibited a greater range of infidelity compared to healthy luminal cells as at least 11% of cells expressed 3 or fewer modules and 59% of cells expressed 6 or more modules. Cancer cells from metastatic samples exhibited additional loss with 30% of cells expressing 2 or fewer luminal modules. **e**. Different patterns of expression for 6 luminal modules, one basal module (L106), and one club module (L56) are present in cells from localized tumor SCG-PCA18. **f.** Euler diagram shows which modules were detected in each cell of localized tumor SCG-PCA18 and illustrates the extent of luminal lineage infidelity within this tumor. **g.** PCA was applied to the luminal modules to quantify the extent of luminal infidelity in each cell. Principal Component 1 (PC1) was averaged across cells to derive a single infidelity score per patient. The infidelity score was significantly lower in localized tumors with Gleason 10 compared to other Gleason scores. **h.** The infidelity score was also significantly lower in patients with higher stage T3 disease compared to T1 and T2 stages. **I.** CaDrA was used to identify combinations of mutations that are maximally associated with module scores that were projected in the TCGA-PRAD cohort. Module L72(*KLK4*) scores were enriched in tumors harboring combinations of *SPOP* mutations, *SPOP* deletions, *CHD1* deletions, or *APC* deletions.

### Infidelity of luminal modules across localized and metastatic PCa

Lineage infidelity within the context of lineage plasticity is characterized by cancer cells losing their original cellular identity and adopting a more undifferentiated state or partial characteristics of a different cell type^35^. We sought to systematically describe the extent of infidelity of the luminal lineage modules within and across PCa samples. Several genes serve as biomarkers for PCa, many of which are associated with the luminal epithelial lineage of the prostate, including *KLK3, AR, NKX3-1, TMPRSS2, KLK4, MSMB, FOLH1,* and *PSCA*^36–43^. Each marker was found in a distinct module including L15, L18, L20, L24, L60, L64, L72, and L97, each of which had distinct expression pattern across cells (**Figure 2c**). The levels of these modules were also quantified in normal luminal cells from three healthy patients^44^. 37%, 22%, and 5% of normal luminal cells expressed 6, 7 and 8 modules, respectively (**Figure 2d)**. Cancer cells from localized tumors exhibited a greater range of infidelity compared to healthy luminal cells as at least 11% of cells expressed 3 or fewer modules and 59% of cells expressed 6 or more modules. Cancer cells from metastatic samples exhibited additional loss with 30% of cells expressing 2 or fewer luminal modules. Luminal infidelity was also a major source of intra-tumoral heterogeneity as expression of the luminal modules varied extensively between cells within many tumors. For example, many tumor cells in SCG-PCA18, a localized stage 2 patient with a Gleason score of 3+3, expressed one of the modules L60 (*NKX3-1*) (85%), L64 (*TMPRSS2*) (94%), and L72 (*KLK4*)(82%). However, only 72% of tumor cells highly expressed all three modules. A subset of cells also expressed other luminal modules or modules for other cell types such as club L56 (PIGR) or basal L106 (*TP63*) demonstrating even more complex infidelity **(Figure 2e,f**). To further investigate the clinical implications of this heterogeneity within localized disease, we quantified the degree of luminal infidelity per patient by calculating the average number of expressed luminal modules per cell as well as average PC1 of all luminal modules per cell, representing a per patient luminal module variation level. This infidelity score was significantly lower in localized tumors with Gleason 10 compared to other Gleason scores (**Figure 2g**). This score was also lower in patients with higher stage disease (**Figure 2h**). Despite these associations, more variation in lineage infidelity was also observed within Gleason and stages groups than between groups raising the possibility that scores of infidelities measured at the single cell level can add additional source for prognostication.

### Somatic drivers of luminal module expression

To determine if somatic alterations are potentially responsible for extensive variation in luminal module expression, we scored each gene module in the TCGA-PRAD bulk RNA-seq data and used CaDrA to determine which combination of mutations or copy number alterations are maximally associated with each module score. Higher levels of the module L72 (*KLK4*) score were enriched in tumors harboring combinations of *SPOP* mutations, *SPOP* deletions, *CHD1* deletions, or *APC* deletions (**Figure 2e**). Co-occurring *SPOP* and *CHD1* alterations have been shown to modulate AR signaling and DNA damage response^45^ and co-occurring *APC* and *SPOP* mutations have also been associated with worse clinical outcomes^46^. Thus, the genes in module L72 likely represent the downstream transcriptional consequences of these alterations and mediate their effect on cellular phenotypes. Besides *KLK4*, other genes in this module such as *AMACR, PART1*, and *SPDEF* are all known to be regulated by AR signaling^47–49^. Some of these alterations were also associated with other gene modules, including an ECM remodeling module L114 (*COL2A1, COL5A1*) and a receptor tyrosine kinase (RTK) signaling module L110 (*ITGAV, ERBB2, GNG12*) showing that these genomic alterations can affect luminal lineage, ECM, and RTK pathways simultaneously (**Supp Figure 2b**). Beyond L72, no other luminal modules were significantly associated with somatic genomic alterations suggesting other mechanisms may be responsible for variability in expression across cells and samples.

### Clinical and biological factors associated with PSA expression

The module L20 contained *KLK2* and *KLK3* which are genes that encode for prostate-specific antigen (PSA), a widely used blood-based marker for PCa prognosis. Interestingly, when examining the distribution of expression for this module across cells, we observed four distinct peaks, which we subsequently named Kallikrein-Depleted, Kallikrein-Downregulated, Kallikrein-Baseline, and Kallikrein-Amplified (**Figure 3a**). Localized PCa samples tended to contain a heterogeneous composition of cells from all four subgroups while CRPC and mCRPC patients predominantly consisted of cells from kallikrein-depleted or kallikrein-amplified subgroups (**Supp Figure 3a**). The proportion of Kallikrein-Downregulated cells was significantly higher in patients who had radiotherapy before prostatectomy (sccomp; FDR < 0.05; **Figure 3b, Supp Figure 3b**) suggesting that a decrease in expression within cells can be driven by external treatment.

**Figure 3.**
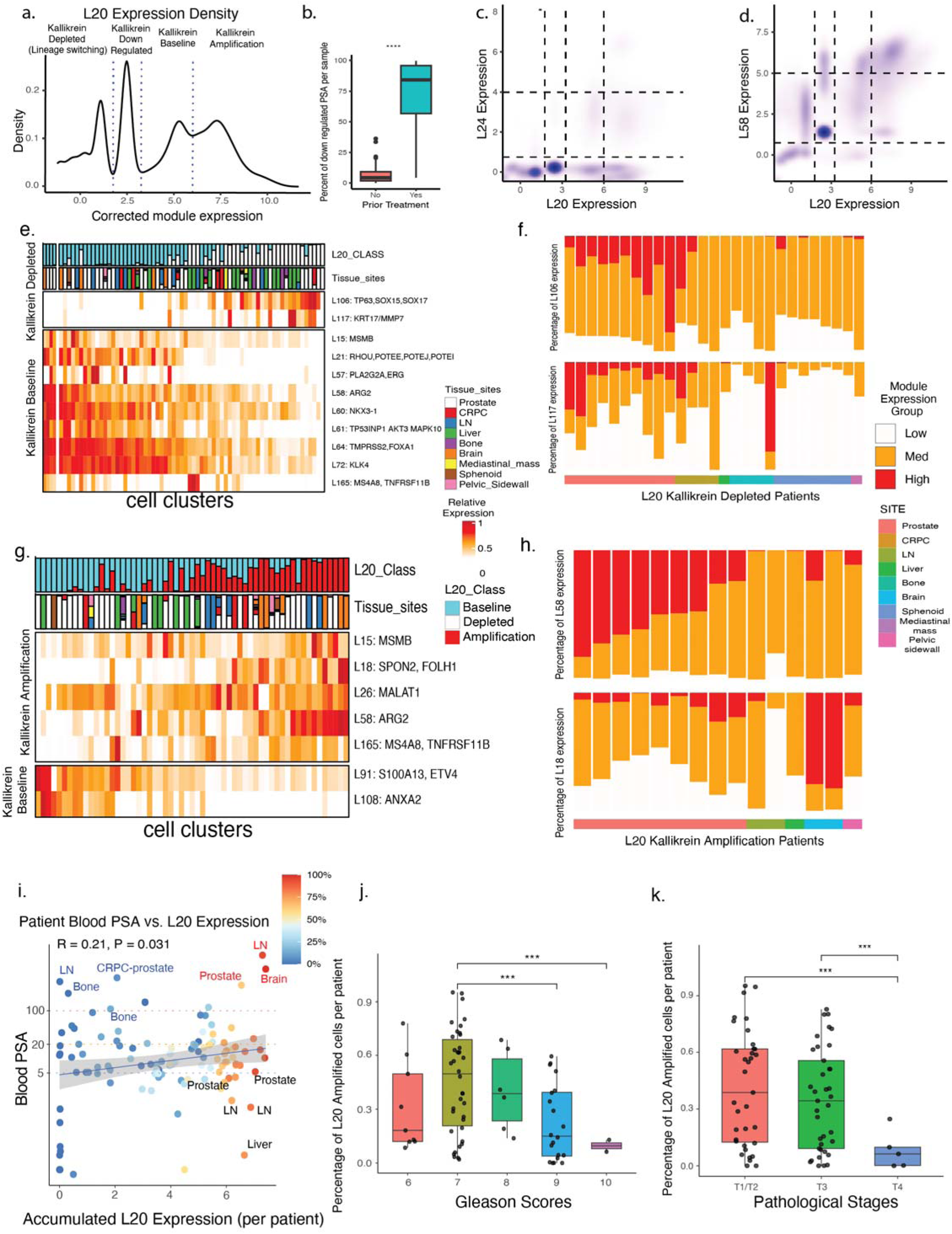
Classification of Kallikrein-related module expression uncovers AR-independent PSA amplification. **a.** Density plot of L20(KLK2/3) module expression was used to divide cells into four groups including Depleted, Down-regulated, Baseline and Amplified. **b.** The percentage of cells classified as “Down-regulated” was significantly lower in localized PCa patients who received radiotherapy prior to prostatectomy (sccomp; FDR < 0.05). **c-d.** 2D contour plot showing L20 expression versus module L24(AR), which contained AR, and L58(ARG2) which was enriched for AR signaling genes. Limited correlation with the AR module suggests that these subgroups are not driven by differences in AR expression. Stronger correlation with the AR responsive module L58 suggests that AR signaling is still active in all kallikrein groups. **e.** Differential module expression associated with L20- depleted cells. **f.** Differential module expression associated with L20-amplification cells. **g.** Composition of L117(KRT17, MMP7) and L106(TP63) expression subtypes in patients with >75% Kallikrein-Depleted cells. **h.** Composition of L18(FOLH1) and L58(ARG2) expression subtypes in patients with >75% Kallikrein-Amplified cells. **i.** Association between patient serum PSA levels and accumulated module L20 expression per sample which is defined as the percentage of Kallikrein-Down-Regulated cells * 2.5 + the percentage of Kallikrein-Baseline cells * 5 + the percentage Kallikrein-Amplified cells * 7.5. Points are colored by the percentage of Kallikrein-Amplified cells. **j**. Association between the percentage of Kallikrein-Amplified cells per sample and Gleason score. **k.** Association between the percentage of Kallikrein-Amplified cells per sample and tumor stage.

As no other clinical variables were associated with Kallikrein-Depleted or Kallikrein-Amplified cells, we sought to determine if other modules were associated with these subgroups of cells. They were likely not driven by transcriptional differences of AR itself as module L24 (AR) was not strongly correlated with module L20 expression (Pearson R = 0.41, **Figure 3c**). However, higher correlation was AR_Signaling module L58 (ARG2) suggesting AR signaling is still active in all four groups (Pearson R = 0.57, **Figure 3d**). Differential module analysis revealed that the basal lineage module L106 (*TP63*) and a mixed basal-luminal lineage module L117 (*MMP7* and *KRT17*) had significantly higher expression in Kallikrein-Depleted cells compared to all other cells (FDR < 0.01; Log2_FC > 0.5) while an ECM associated module L108 (*ANXA2*) was significantly lower (FDR < 0.01; Log2_FC < -0.50; **Figure 3e**). 10 localized, 4 localized CRPC, and 13 metastatic samples had at least 75% of kallikrein-depleted cells. The 10 localized samples had a mix of cells that were high or intermediate for L106 while the 13 metastatic samples mostly harbored cells with intermediate expression of L106. Conversely, module L117 generally displayed low to intermediate expression across metastatic samples except for one brain metastatic sample (**Figure 3g**, **Supp Figure 3c**). These observations suggest that intermediate and high expression of module L106 potentially coupled with the presence of module L117 marks a transition between luminal and basal cell states with a concurrent depletion of kallikrein module.

The kallikrein-amplified cells were not likely driven by transcriptional up-regulation of AR itself as module L24:AR was not strongly differently expressed between kallikrein-amplified and kallikrein-baseline cells (Log2_FC = 0.1). Rather, cells in the kallikrein-amplified group had significantly higher expression of the AR signaling module L58 (*ARG2*) as well as luminal modules L15 and L18 **(Figure 3g, Supp Figure 3d)**. 9 localized, no CRPC, and 6 metastatic samples had at least 75% of kallikrein-amplified cells (**Figure 3h**). The 9 localized samples had higher fractions of L58-Intermediate or L58-High cells suggesting that AR signaling is maintained or increased via other mechanisms beyond mRNA expression of the AR gene itself. L18-Intermediate cells were found across most tumors with the exception of two brain metastatic samples, which exhibited a high fraction of L18-High cells. Module L58 displayed consistently intermediate expression across metastatic samples compared to localized PCa samples which had higher fractions of L58-High cells. These findings suggest that module L58 may function as an initial driver facilitating PSA overamplification, while high levels of module L18 could be implicated in subsequent metastatic stages.

We next examined the relationship between L20 (KLK2/3) module expression and serum PSA. Across 109 samples with PSA data, we observed a significantly positive correlation between the accumulated kallikrein module expression per sample and serum PSA levels obtained from the patient at the nearest timepoint (R = 0.21, p = 0.031, **Figure 3i**). Despite the overall correlation, there were notable exceptions of samples that had discrepancies between serum PSA and tumor cell gene expression. In 2 bone metastases, 1 LN metastasis, and 1 CRPC prostate sample, the accumulated kallikrein module expression was relatively low despite extremely high levels of serum PSA (>100). This pattern may reflect contributions from non-malignant sources such as benign prostatic hyperplasia (BPH), the primary tumor, or other metastatic sites that were not captured. Conversely, 2 LN and 1 liver metastatic samples displayed high accumulated kallikrein module expression without a corresponding increase in serum PSA, suggesting a decoupling between kallikrein transcription and PSA release in these samples. When examining pathological and clinical phenotypes, serum PSA was higher in intermediate risk tumors with Gleason 7 and 8 tumors having significantly higher PSA compared to Gleason 6 tumors (**Supp Figure 3h**). In contrast, the loss of Kallikrein-Amplified cells was more strongly associated with more severe phenotypes. A significantly lower proportion of Kallikrein-Amplified cells were observed in tumors with Gleason score 9 or 10 compared to Gleason 7 tumors and in patients with T4 disease compared to T1/T2 (**Figure 3j,k, Supp Figure 3i-m**). These findings indicate that while the cellular heterogeneity of kallikrein expression contributes to variability in serum PSA, it does not fully account for the highly discordant PSA levels observed in some patients with advanced grade and stage disease.

### The landscape of gene modules associated with PCa metastasis

Cancer cells undergo extensive reprogramming to acquire traits that facilitate metastasis to distant sites. To investigate the molecular basis of metastasis, we identified gene modules with significantly altered expression in metastatic PCa compared to localized disease. Two modules related to the cell cycle, L12, and L156, had significantly higher expression in metastatic disease across all sites (FDR < 0.05; |Log2 FC > 1.3|; **Figure 4a,b**). L12 was enriched with genes from the G2/M phase such as *MKI67*, *TOP2A*, and *CCNB1/2*^50,51^, while L156 was enriched for genes in G1/S phase including those involved in spindle-assembly and INK4 checkpoint genes such as *UBE2S, TUBA1A* and *CDKN2A/C/D* (**Supp Figure 4a-c**)^52–54^. We annotated cells as “Inactive”, “Intermediate”, and “Active” states corresponding to low, moderate, and high expression for each module (**Figure 4c**). As expected, metastatic samples contained a larger fraction of Active cells on average compared to localized tumors, reflecting higher rates of proliferating cells in these more aggressive tumors. In average, 25% and 17% of cell were Active for L12 and L156 per sample among metastatic tumors, respectively compared to only 4% for both L12 and L156 per sample among localized tumors (**Figure 4d**). Interestingly, metastatic samples were also dominated by cells in the Intermediate state of either module. Specifically, 56% of metastatic cancer cells per sample were Intermediate for module L12 and 70% were Intermediate for module L156 which was significantly higher than localized disease (44% for L12 and 46% for L156). These findings indicate that many cells from metastatic samples and some from localized disease may occupy a transcriptional state poised for proliferation even when they are not actively undergoing mitosis.

**Figure 4.**
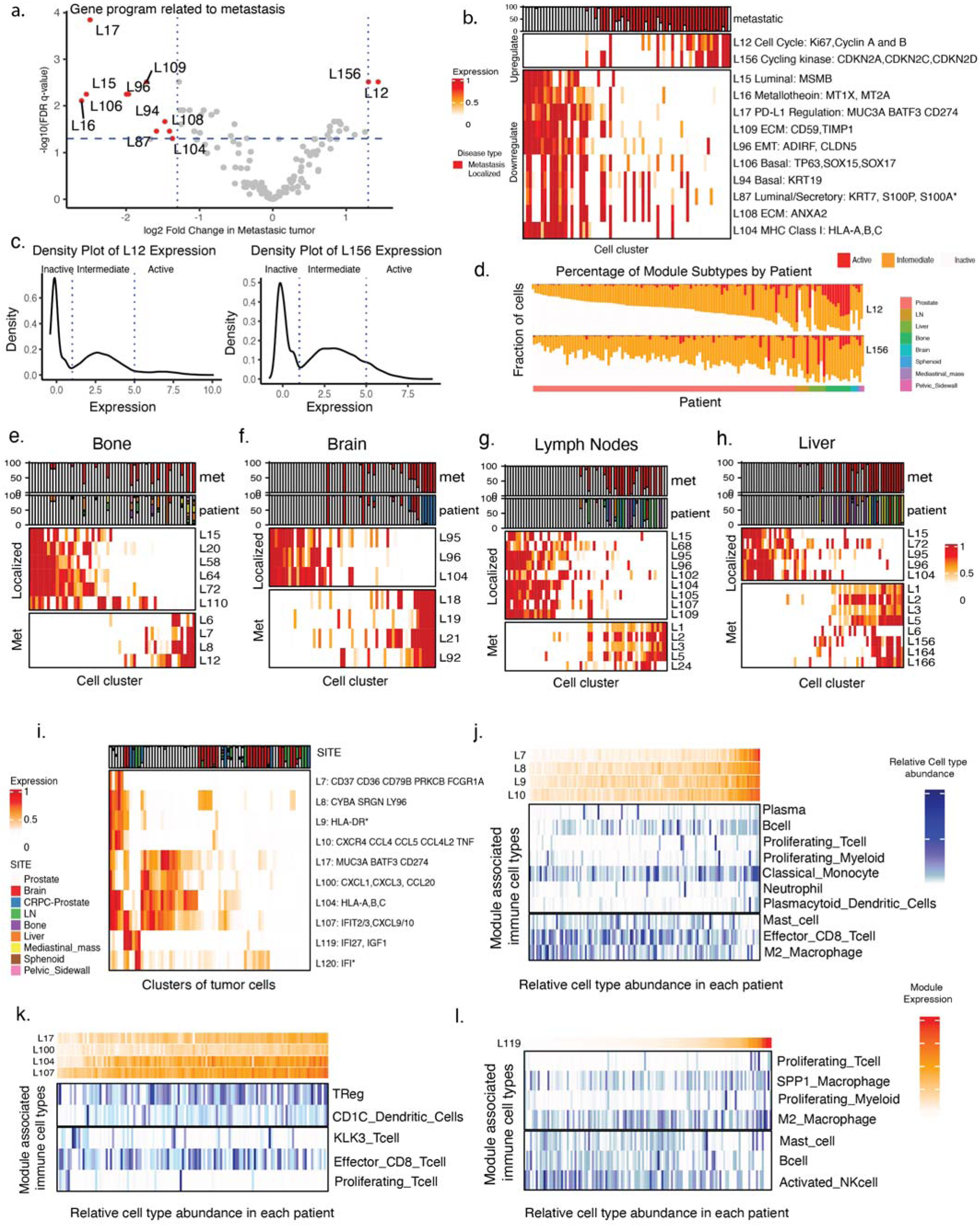
Landscape of metastasis- and immune signaling-associated modules. **a.** Linear mixed effect modeling (LME) was used to identify modules associated with all metastatic samples regardless of organ site while controlling for study, tumor, and cluster as nested random effects. A volcano plot shows the Log2 Fold Change (i.e. the model coefficient for metastasis) versus the -Log10 FDR q-value. **b.** Heatmap of modules significantly associated with metastasis (FDR < 0.05; |Log2 Fold Change| > 1.3). **c.** Density plot of cell cycle modules L12(Ki67) and L156(CDKN2A/C/D). Based on these densities, cells were classified as “Inactive”, “Intermediate”, or “Active” for each module. **d.** Distributions of L12 and L156 cell groups per patient. Metastatic samples had higher proportions of “Active” cells although “Intermediate” cells for both modules could be also observed in both metastatic and localized samples. **e-h**. LMEs were used to identify modules associated with each individual organ site compared to localized tumors. Each heatmap shows modules associated with a specific organ site (p-value < 0.05). **i.** Heatmap shows the expression of modules annotated as “immune” or “signaling” across tumor cell clusters. 3 broad axes of signaling could be observed in cells from localized disease including 1) cells with higher levels of L7, L8, L9, and L10; 2) cells with higher levels of L17, L100, L104, and L107; and 3) cells with higher levels of L119. **j-l.** Immune cell type abundances were associated with immune signaling modules expression using sccomp. Each heatmap shows the abundance of immune cell populations that were significantly correlated with the per patient the modules scores in each axis (FDR < 0.05).

Several gene modules were significantly downregulated in metastatic prostate cancer compared to localized disease. These included multiple lineage-associated programs including L15 (*MSMB*) and L87 (*KRT7, S100P, S100A14/16, SERPINB1*), L106 (*TP63, SOX15/17*) and L94 (*KRT19*)^55^. In addition, we observed a reduction in ECM remodeling–related modules, including L96 (*ADIRF, CLDN5*)^56^, L109 (*COL6A1/2*), and L108 (*ANXA2*)^57^, relative to localized cancer cells. Collectively, these decreases suggest that metastatic progression is accompanied by a loss of prostate lineage-defining programs and a shift towards a more plastic cellular state with decreased cell adhesion. We also noted a marked downregulation of the metallothionein module L16 (*MT1A/E/F/G/H/M/X, MT2A*), indicating a disruption of prostatic zinc homeostasis, which has previously been implicated in disease progression^58^.

We also performed additional differential analyses to identify specific modules associated with each organ metastatic site compared to localized disease (**Figure 4e-h**). Several modules were uniquely enriched in specific sites and displayed functional characteristics of their metastatic microenvironment. For example, module L8 had significantly higher expression in bone metastasis and contained *LGALS1* (Galectin-1, a tractable target in CRPC)^59^ along with hematopoietic/osteoclast regulators *CYBA, SRGN*, and *LY96* (**Figure 4e; Supp Figure 4d-f**). We further validated the elevated module expression of L8 is specific to bone metastasis supported by bulk RNA-seq dataset (**Supp Figure 4g**). Module L19 was induced in brain metastasis and composed of axon-guidance and neuro-migratory genes *EPHA4*, *EPHB2*, and *MAP1B* (**Figure 4f**; **Supp Figure 4h-j**). Other modules such as L5 were associated with multiple metastatic sites including lymph-node, liver, and mediastinal-mass (**Figure 4g,h**; **Supp Figure 4k-m)**. L5 was enriched for glutamatergic-signaling genes (*GRIN2A, GRIN2B, GRIA3, GRM7*) and stromal-remodeling regulators (*SOX5, TGFB2, FGFR2*) suggesting that excitatory neurotransmission and TGF-β–driven matrix modulation are shared features of metastatic colonization (**Figure 4g,h**; **Supp Figure 4k**). In addition, module L6 contained multiple S100A family members whose expression was significantly elevated in both liver and bone lesions. The liver metastases displayed a dichotomous, patient-specific pattern of L6 where cells from 2 tumors had high proportion, and 3 other tumors had low proportion. In contrast, all the 9 bone metastatic samples showed a uniform pattern of intermediate proportion of L6-expressed cells (**Supp Figure 4l-o)**. Overall, these patterns suggest that metastatic transcriptional states comprise both niche-adaptive modules, which are preferentially deployed to tolerate site-specific stresses and maintain viability in restrictive microenvironments, and core modules that support metastatic growth more broadly. Even among the core modules, site-dependent attenuation or amplification indicates ongoing microenvironmental shaping of otherwise shared tumor-intrinsic programs.

### Heterogeneity of immune and signaling modules

Immune checkpoint blockade therapies have shown limited efficacy in PCa patients demonstrating the need to characterize immune activation and evasion pathways^60^. Several immune-related modules were largely lost in metastatic lesions, including the MHC class I antigen presentation module L104 (*HLA-A/B/C*) and the PD-L1–associated module L17 (*MUC3A, CD274*)^60^. Module L100 (*CXCL1*, *CXCL3, CCL20*) contained several cytokine and immune signaling genes, was also more prevalent among localized disease, and co-occurred with L17 in a subset of localized samples. Two distinct interferon response modules were detected including L120 (*IFI6, IFI44/44L*) which was moderately expressed across many cells and L119 (*IFI27*) which was highly regulated in a smaller subset of cells. Several modules were enriched for genes involved in immune differentiation including L7 and L8 as well as module L9 containing MHC class II genes. L7, L8, L9, and L10 were highly expressed in a smaller subset of cells that largely lacked high expression of L17, L100, L104 and L107, and L119 were expressed in cells that were depleted of both the of groups of modules, suggesting 3 broad immune signaling axes in cells (**Figure 4i**). To further characterize these three axes, we related module expression in tumor cells to the composition of infiltrating immune cell types across all tumor samples. Higher average expression of L7–L10 in tumor cells was associated with an immune-supportive microenvironment enriched for proliferating T cells, myeloid cells, and B cells. In contrast, increased expression of L17/L100/L104/L107 was associated with an immunosuppressive environment characterized by upregulation of Treg and CD1C⁺ populations and a concomitant reduction in CD8⁺ T cells. Finally, L119 expression was associated with a more myeloid-dominated signature, consistent with a myeloid-specific axis of immune modulation. Together, these patterns suggest that module expressions in tumor cells may reflect distinct mechanisms of immune microenvironment remodeling, in which specific transcriptional programs selectively recruit or expand specific immune cell types while limiting the infiltration or activity of others, thereby enabling alternative routes of immune escape in localized disease.

### Landscape of gene modules in cancer cells associated with NEPC

Differentiation to neuroendocrine-like (NE) phenotypes is a mechanism that can lead to castration resistance and metastasis. To further characterize and refine gene expression patterns of NEPC within and across tumors, we identified 11 gene modules that were significantly enriched in tumors clinically annotated at NEPC compared to all other samples (FDR<0.05, |Log Fold Change|>1; **Figure 5a**). Module L166 had the second highest fold change and enriched for markers of canonical NE program including *SCG2*/3. Surprisingly, module L164 was more strongly upregulated than the canonical NE module. L164 contained HES6, a basic helix-loop-helix transcriptional repressor previously implicated in androgen independent growth and castration resistance^61,62^. We considered L164 and L166 as “core” NE modules given that they shared genes with previously established NE signatures such as SCG2/3 and SYP, however they contained other genes not included in previous signatures (**Supp Figure 5a**).

**Figure 5.**
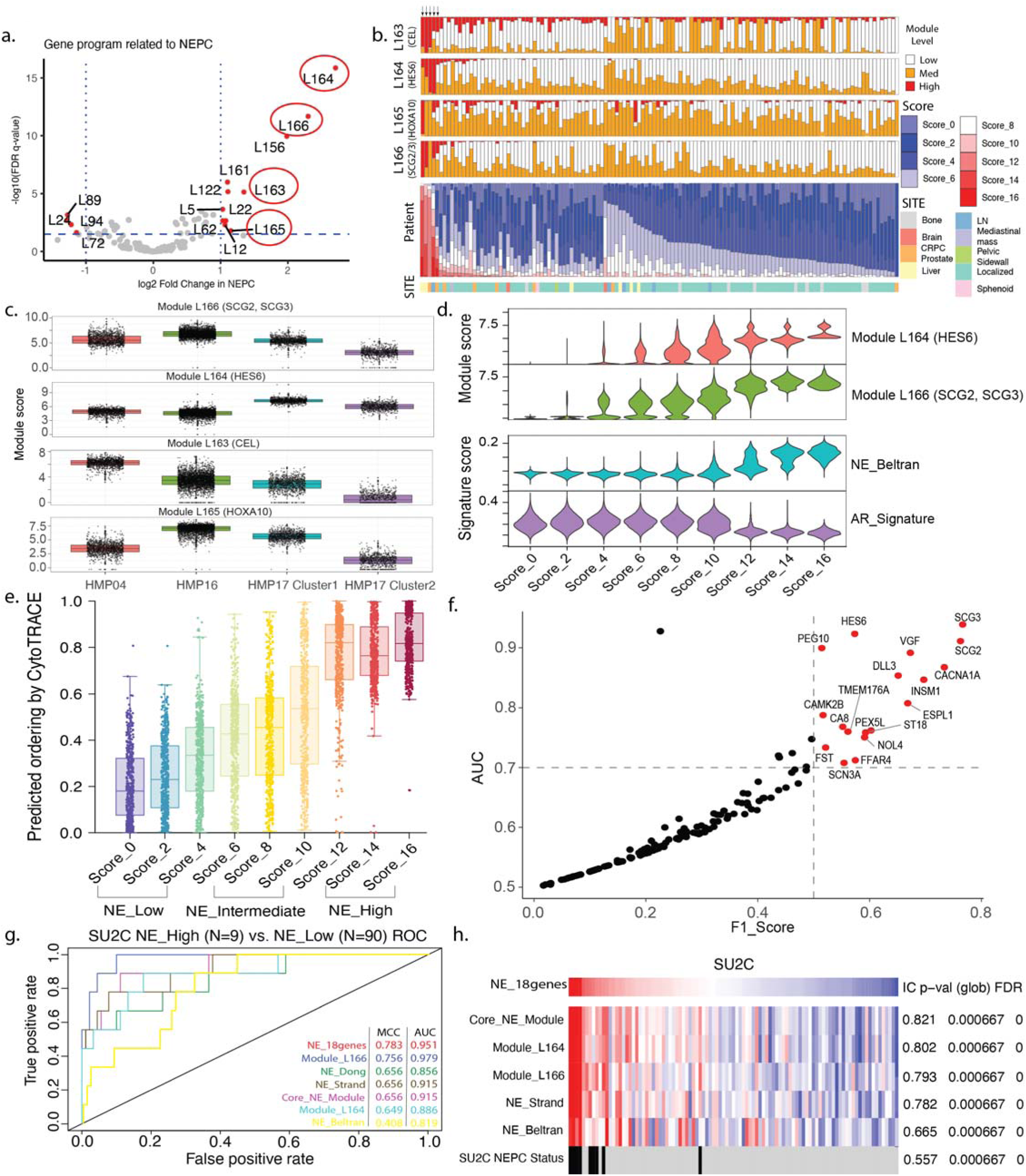
Neuroendocrine-like tumor analysis reveals precursor-like intermediate neuroendocrine cells. **a.** Identification of significantly enriched tumor gene programs in NEPC patients (right, N = 5) compared to non-NEPC patients (left, N = 123). Significantly enriched modules are specifically labeled. The two “core” and two “supporting” NE modules are circled. **b.** Upper panel, distribution of all four NEPC-enriched module scores for all 128 patients; arrows mark the clinically annotated NEPC patients. Lower panel, Stacked barchart of tumor cells for each patient colored by NE scores, sorted by the contribution of high NE scoring tumors. The annotation bar shows the site for each patient. **c.** Box plots of module scores for the four NE modules in three NEPC patients (HMP04, HMP16, HMP17). **d.** Stacked violin plots of the two core NE modules and two established signatures across tumor cells of different NE scores. **e.** CytoTRACE differentiation box plots for all tumor cells colored by the computed differentiation scores NE scores are grouped in three categories. **f.** F1-score and AUC for genes in the core NEPC-enriched modules and the 18 genes with highest prediction accuracy are highlighted in red. **g.** ROC curve of NE-high signature scores (left). AUC and MCC are shown in the plot. **h.** ssGSEA score heatmap of NE-high signature in the SU2C dataset and correlation with other NEPC-enriched tumor modules and NEPC annotation. Information score (IC), association p-values and FDR are shown next to the heatmap. Black bars indicate NEPC patients in the SU2C cohort.

Beyond these two “core” NE modules, we considered L163 and L165 as “supporting” modules that were associated with NEPC status. Module L163 contained CEL, a previously described marker for pancreatic acinar cells^63^, along with IGFBP5, an insulin growth factor. Module L165 containing HOXA10 was also upregulated in the Amplified KLK2/3 expressing cells suggesting that it can promote multiple phenotypes. Beyond these four “core” and “supporting” modules, several other modules had significantly higher expression in NEPC. However, their functional annotations and broader expression across tumor cells suggest that they are less specific to neuroendocrine lineage transformation and instead reflect more general programs associated with advanced disease. These included cell cycle module L12, module L5 which was associated with metastatic samples in general, as well as cyclin kinase module L22, AR pathway associated module L62, cytoskeleton module L122 and cellular membrane remodeling module L161. Notably, although module L156 appeared to be a statistically significant NE associated module, the genes expressed in this module are mostly cell cycle G1/S checkpoint associated genes and thus we determine its association with NEPC is more relevant to cell proliferation rather than the specific neuroendocrine lineage. In general, the large number of samples of this atlas combined with this refined module analysis allowed for a detailed characterization of the various biological programs associated with NEPC transformation at single-cell resolution.

We next sought to characterize the heterogeneity of NE-associated modules within and across tumors. Each cell was categorized as having Off (low), Intermediate, or Active (high) expression for each of the 2 “core” or 2 “supporting” NE modules (**Supp Figure 5b)**. Interestingly, each NEPC tumor exhibited a unique pattern of cells with high levels of these modules (**Figure 5b**). For example, when examining the core NE modules among NEPC patients, cells from HMP16 harbored higher levels of L166, while cells from HMP17 and HMP04 contained cells with higher levels of L164 (**Figure 5c**, **Supp Figure 5c**). Similarly, among the supporting modules, cells from HMP04 contained high levels of L163 and low levels of L165, cells from HMP16 contained intermediate levels of L163 and high levels of L165, while cells from HMP17 contained lower levels of L163 and L165. These patterns demonstrate cells from NEPC are not monolithic but can be defined by different combinations of NE-associated gene modules.

### Heterogeneous expression of NEPC-associated modules observed in localized disease

Expression of NE markers has been reported in cells from localized PCa suggesting that the transition to a NE phenotype can start before clinical progression^64,65^. However, these studies have primarily relied on a few protein markers or bulk RNA-seq data. In this atlas, we observed some cells with high or intermediate levels of core NE modules in samples not clinically annotated as NEPC. Specifically, 1 localized sample had greater than 5% of cells with high levels of the canonical NE module L164 while 54 samples had at least 25% of cells with intermediate levels of L164 (**Figure 5b**). Similarly, 6 localized samples had at least 5% of cells with high levels of L166 while 47 localized tumors had at least 25% of cells with intermediate levels of L166 demonstrating that some degree of expression of these lineage programs can be broadly detected at lower levels in subsets of cells.

To define the cumulative extent of NE transformation in each cell, we developed a NE scoring system which is sum of scores from the two NE core and two supporting modules (**Methods**). The overall NE score for each cell was computed as the sum of all four module scores and ranged from 0 to 16. Each cell was then classified as NE low (0-4), intermediate (6-10), or high (12-16) based on the overall NE score (**Figure 5b, Supp Figure 5d**). As expected, the expression for modules L164 and L166 displayed a stepwise progressive increase across the nine resulting NE score groups, consistent with a continuum of NE differentiation rather than large state transitions (**Figure 5d**). The coordinated increase in NE “intermediate” cells was not observed for previously published NE signatures which only had higher levels in the “high” NE scoring cells (**Supp Figure 5e)**. To validate the continuum of differentiation, we used CytoTRACE to estimate the degree of differentiation in each cell. CytoTRACE scores also displayed an increasing stepwise trajectory from “low” to “high” NE-scoring cells (**Figure 5e**), supporting the idea that cells with intermediate expression of NE modules may represent transitional or primed differentiation states. We also performed differential expression analysis followed by gene set enrichment between the three major NE score categories to identify additional pathways. NE low cells were enriched in androgen response, TGF beta and interferon gamma signaling, NE intermediate cells were enriched for G2M and E2F signatures, while NE high cells were enriched for DNA repair pathways (**Supp Figure 5f**). The increasing degree of differentiation and distinct biological pathways in cells with intermediate NE states supports a model in which molecular plasticity begins to occur before occult histological transformation.

### Improved signatures for prediction of NEPC

Previously established neuroendocrine signatures from Beltran et al and Dong et al^15,27^ were enriched in cells from clusters 8 and 10 (**Figure 1f**, **Supp Figure 5e**). Despite enrichment of both signatures in cells from NEPC patients, they displayed heterogeneous and discordant expression across the remaining samples. To derive a more precise predictor of NE-like states, we evaluated genes within Modules L164 and L166 by calculating their AUC, PR-AUC, F1 scores, and MCC for classifying NE-high versus other tumor cells (**Figure 5f, Supp Figure 5g, Supp Table 3**). This analysis yielded an 18-gene signature with overall strong predictive performance in our atlas (FDR < 0.05, AUC >= 0.7, F1 score >= 0.5). To validate the predictive capability of the core NE modules and the refined 18-gene signature in an independent bulk RNA-seq dataset, we scored each signature in the SU2C bulk RNA-seq dataset which contained 9 clinically annotated NEPC/CRPC samples. The 18-gene signature achieved the highest MCC (0.783) and the second highest AUC (0.951) behind the full set of genes from module L166 for distinguishing NEPC from non-NEPC samples (**Figure 5g**). The ROC curves (**Figure 5g**) illustrated overall classification performance, the heatmap (**Figure 5h**) provided an intuitive view of individual NEPC-positive patients being distinctly clustered and scored highly for the 18-gene NE signature and L164/166 modules, supporting their accuracies and utilities in characterizing the NEPC phenotype (**Supp Figure 5h)**. Importantly, the 18-gene NE signature and module L164 had better performance than previously derived NE signatures demonstrating its potential as a robust marker set for identifying NE-like phenotypes in prostate cancer in bulk RNA-seq datasets. Overall, these findings reveal substantial heterogeneity in NE programs across tumors and nominate a concise gene signature that can facilitate classification of NEPC.

### Distinct stromal and immune cell states characterize localized and CRPC prostate cancer

To complement our tumor cell module analysis, we performed unsupervised clustering of stromal and immune compartments to investigate their cellular diversity across disease states. Stromal clustering identified distinct populations including ECM fibroblasts, smooth muscle cells, IFN-responsive fibroblasts, and MSC stromal cells (**Supp Figure 6a**), and each cluster was characterized by a group of differentially expressed genes (**Supp Figure 6b**). Analysis of stromal composition across localized PCa samples, CRPC prostate samples, and mCRPC samples showed that mCRPC and CRPC samples harbored significantly more ECM fibroblasts (**Supp Figure 6c**). Pathway enrichment analysis showed differential activation of hallmark pathways, including upregulation of interferon signaling in IFN fibroblasts and epithelial-mesenchymal transition pathways in ECM fibroblasts (**Supp Figure 6d**). Given the enrichment of ECM remodeling and fibroblast activation genes, we next assessed whether these changes reflected shifts in cancer-associated fibroblast (CAF) states using the CAF core matrisome signature. CAF core matrisome signature scores were significantly enriched in CRPC prostate and mCRPC samples compared to localized PCas (**Supp Figure 6e**), supporting increased fibroblast activation and ECM deposition in advanced disease. Gene set enrichment analysis further demonstrated upregulation of matrisome pathways and ECM organization in both CRPC prostate and mCRPC stromal compartments (**Supp Figure 6f,g**).

**Figure 6.**
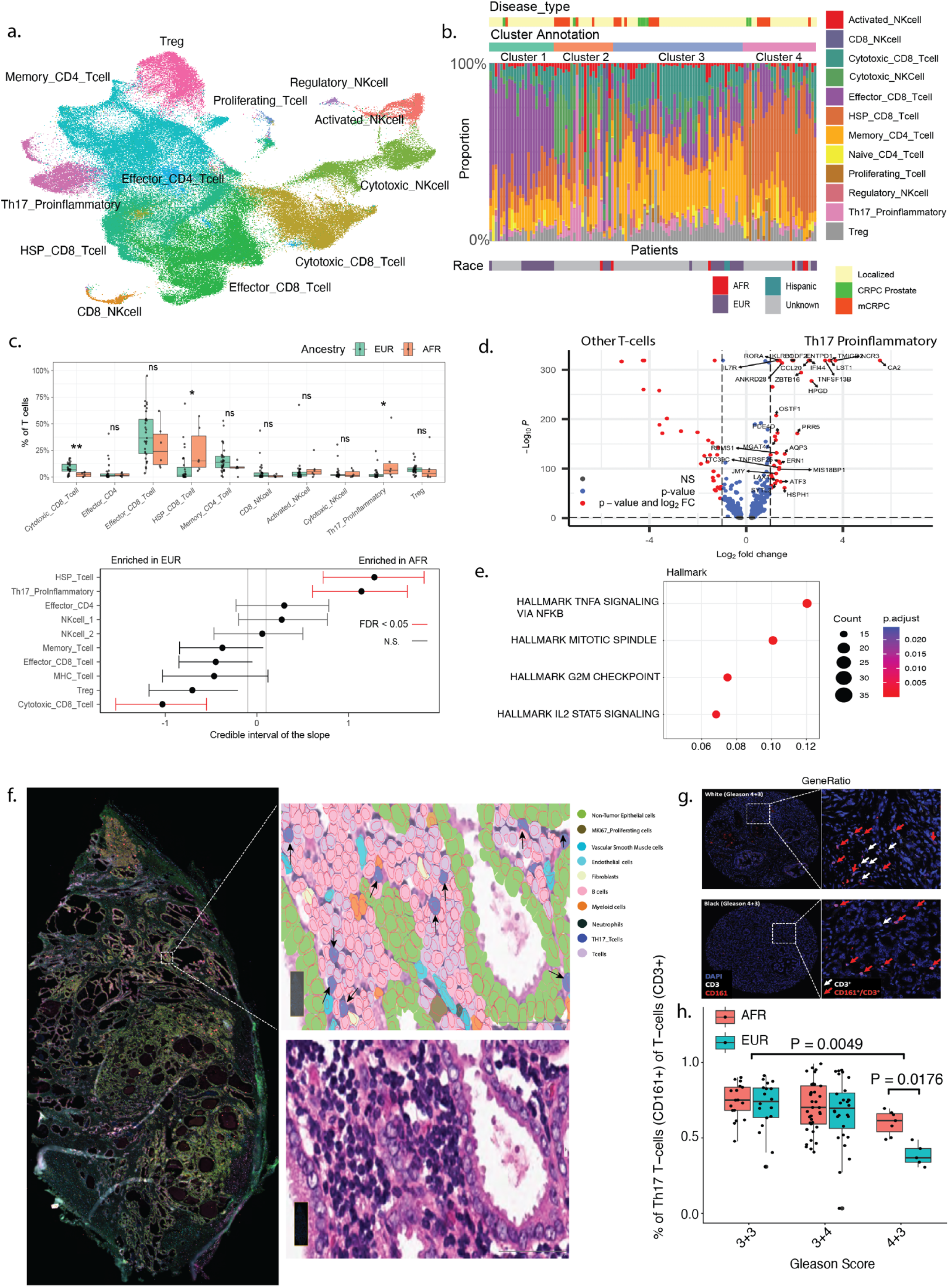
TME analysis reveals enrichment of Th17 T-cells and HSP T-cells in African American PCa. **a.** UMAP visualization of T-cell subclusters identified from the integrated single-cell atlas. Cell types are labeled on the UMAP. **b.** Patient-level clustering based on T-cell subtype composition identifies four patient clusters with distinct immune profiles. Stacked bar plots show the relative proportion of T-cell subtypes per patient. Disease type and self-reported races are shown above and below the heatmap. Cluster 4 is characterized by enrichment of HSP_CD8_Tcells and contains a higher proportion of CRPC and mCRPC samples. **c.** Comparison of T-cell subtype proportions between European American (EUR) and African American (AFR) patients. Top: boxplots showing percentage of each T-cell subtype among total - cells. Bottom: compositional modeling using sccomp showing credible intervals of ancestry-associated differences. Significantly enriched populations are labeled in red (FDR < 0.05). **d.** Differential gene expression analysis comparing Th17 proinflammatory T-cells to other T-cell populations. Volcano plot highlighting canonical Th17 markers including *RORC*, *CCL20*, *GPR65*, *KLRB1*, and *CXCR6*. **e.** Gene set enrichment analysis (GSEA) result of significantly enriched pathways of genes enriched in Th17 T-cells from AFR patients. **f.** Spatial transcriptomic profiling using the Xenium 5K Prime platform in prostate cancer tissue from an AFR patient. Th17 cells are marked by KLRB1 staining. Right panels show cell type annotation and histological context in the zoomed-in region. **g.** Representative mIF images of PCa tissue TMAs (DAPI = nuclei, CD3 = T-cells, CD161 = Th17 marker) from EUR (top) and AFR (bottom) patients with Gleason 4 + 3 tumors. White arrows indicate CD3⁺ cells; red arrows indicate CD161⁺ CD3⁺ (Th17) T-cells. **h.** Boxplots showing the percentage of Th17 T-cells in all T-cells across Gleason grades (3 + 3, 3 + 4, 4 + 3) in AFR (red) and EUR (blue) patients. Statistical significance is indicated.

In the immune compartment, clustering revealed major T cell, NK cell, and myeloid populations after excluding potential contamination to improve robustness of immune. Myeloid cell clustering identified classical monocytes, CD16⁺ non-classical monocytes, neutrophils, conventional dendritic cells (cDC1 and cDC2), plasmacytoid dendritic cells (pDCs), TAM-like macrophages, and proliferating myeloid precursors (**Supp Figure 6h**). Using compositional analysis with sccomp, we identified localized PCa to CRPC shifts in myeloid populations, with proliferating myeloid cells and plasmacytoid dendritic cells significantly enriched in CRPC, while classical monocytes and neutrophils were more abundant in localized prostate cancer (**Supp Figure 6i**). These analyses provide an overview of stromal and immune heterogeneity within the tumor microenvironment.

### Enrichment of Th17 T-cells in African American PCa

We performed a sub-clustering analysis of T-cells and identified regulatory T-cells, effector and memory CD4⁺ T-cells, effector and cytotoxic CD8⁺ T-cells, HSP-high CD8⁺ T-cells, Th17 pro-inflammatory T-cell cluster^66,67^ (marked by *RORA*, *IFI44*, and *CA2*), and proliferating T-cells, while NK cell clusters comprised cytotoxic, regulatory, activated, and CD8⁺ NK cells (**Figure 6a**). To investigate patient-level T-cell heterogeneity, we next clustered patients based on their cell type composition. This analysis identified four patient groups with distinct immune profiles (**Figure 6b**). Notably, one patient cluster (Cluster 4) exhibited a marked enrichment of HSP_CD8_Tcells and contained a higher proportion of CRPC and mCRPC samples compared with other clusters. Cells within this population showed elevated expression of stress-response genes including *HSPA6*, *HSPA1A*, *DNAJB4*, and *HSPA1B*, while maintaining expression of canonical T-cell markers such as *CD3D* (**Supp Figure 7a**), consistent with a stress-associated CD8⁺ T-cell state.

**Figure 7.**
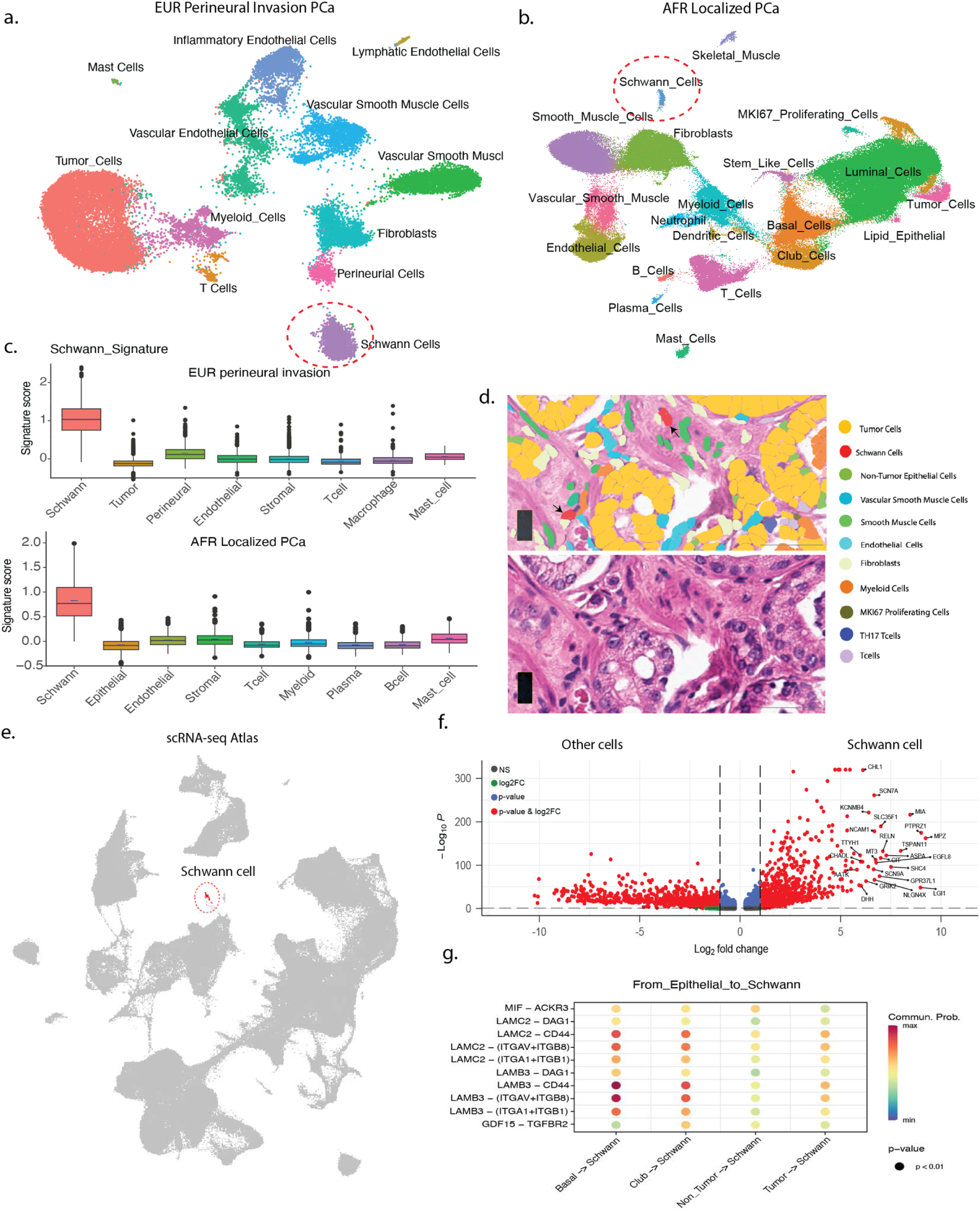
Spatial transcriptomic identification of Schwann cells in localized prostate cancer. **a–b.** UMAP showing cell type annotations from Xenium 5K spatial transcriptomic profiling of one European American perineural invasion (panel a) and one African American localized (panel b) prostate cancer tissue sample. Schwann cells are specifically identified and highlighted in both samples. **c.** Box plot showing the Schwann cell signature score for all major cell types in both Xenium samples. **d.** Upper panel: H&E image of prostate tissue with corresponding Xenium segmentation and cell type annotation overlay. Schwann cells are shown in red and are located adjacent to tumor glands (orange) and endothelial cells (green). Black arrows indicate nerves or perineural regions. Lower panel: Corresponding H&E section showing tumor and nerve structures. **e.** Integrated UMAP of the atlas highlighting the localization of Schwann cells (red). **f.** Volcano plot showing differentially expressed genes in Schwann cells compared to all other cell types. Canonical Schwann cell markers are highlighted. **g.** CellPhoneDB ligand-receptor interaction analysis between Schwann cells and epithelial populations (basal, club, luminal, and tumor cells). Shared interactions, such as LAMB3–ITGA and LAMC2–ITGA, with dot size indicating interaction significance and color scale showing the communication probability.

Interestingly, we also noticed that 3 of the 6 AFR patients with available ancestry information clustered within Cluster 4 (**Figure 6b**, **Supp Table 1**). Ancestry admixture analysis supported the self-reported race information for these samples (**Supp Table 4**). This observation led us to perform a systematic comparison of T-cell subpopulation composition between AFR and EUR patients. Previous bulk transcriptomic studies have reported enrichment of inflammatory pathways in AFR prostate cancer patients^68,69^. However, few AFR PCa samples have been characterized at single-cell resolution. Among the 98 localized PCa samples in our single-cell atlas, 31 patients were European American (EUR) and 6 were African American (AFR) according to self-reported race and ancestry analysis (**Supp Table 1, Supp Table 4**).

Comparison of T-cell subpopulation proportions revealed that Th17 and HSP⁺ T-cell clusters were significantly enriched in AFR patients (T-test, FDR < 0.05; sccomp, FDR < 0.05; **Figure 6c)**, while cytotoxic CD8⁺ T cells were present at lower proportions (**Figure 6c**). Specifically, Th17 T-cells represented 12.3% of all T-cells in the AFR group compared with 4.0% in the EUR group, while HSP⁺ T-cells accounted for an average of 28.7% of T-cells in AFR patients versus 8.4% in EUR patients (**Figure 6c, Supp Figure 7b**). Differential expression analysis of the Th17 population revealed increased expression of canonical Th17 markers including *RORC* (encoding RORγt), *CCL20* (a key Th17 chemokine), and *GPR65*, together with surface markers such as *KLRB1* (CD161) and *CXCR6* (**Figure 6d**). Gene set enrichment analysis showed these genes were significantly associated with IL2–STAT5 signaling, TNFα signaling via NFκB, and inflammatory response pathways (**Supp Figure 7c**), consistent with the pro-inflammatory nature of Th17 cells.

This Th17 T-cell abundance was also found to be positively associated with other T-cell subsets, including effector CD4, memory T-cells, Tregs, and cytotoxic CD8 T-cells, and most strongly anti-correlated with ECM and activated fibroblasts (**Supp Figure 7d**). This indicated that Th17 T-cells expand concordantly with other T-cell populations, while their abundance is reduced with increasing fibroblast activity, which suggested a balance between immune cell infiltration and stromal activation within the TME.

Among Th17 T-cells, those from AFR PCa showed enrichment of TNFA signaling via NFkB (**Figure 6e**), marked by NFκB regulators including NFKBIZ and NFKBIE, as well as TNFAIP3, TRAF3, and TNIP2 (**Supp Figure 7e**). To determine the spatial localization of Th17 cells in PCa tissue, we performed spatial transcriptomic sequencing using the Xenium 5K prime platform on a prostate cancer tissue from an AFR patient. CD3D+ and KLRB1+ T cells, consistent with a Th17 identity, colocalized with a large aggregate of B cells and other immune cells resembling tertiary lymphoid structures (**Figure 6f**). To further characterize the heterogeneity of Th17 T-cells in prostate cancer, we performed multiplex immunofluorescent (mIF) staining for CD3 and CD161 (KLRB1) in two prostate cancer tissue microarrays (TMAs) comprising cores from 29 AFR and 27 EUR patients. Th17 cells ranged from undetectable to >100 cells per core and occupied both stromal and epithelial niches. Within the TME, Th17 T-cells were frequently embedded in acute or chronic inflammatory infiltrates and in some cases, within formations resembling tertiary lymphoid structures (**Figure 6g**). Notably, Th17 T-cells were also found in close proximity to benign epithelial or malignant glands, underscoring a potential for direct spatial crosstalk with epithelial cells. The enrichment of these pro-inflammatory cells in AFR PCa patients may contribute to the inflammatory nature of prostate cancer that has been described.

To validate the enrichment of Th17 T-cells observed in our single-cell analysis, we then quantified Th17 T cells in the two TMAs. Representative images demonstrated a higher percentage of CD161⁺ CD3⁺ T-cells within tumor regions from AFR Gleason 4+3 patients compared with EUR Gleason 4+3 patients (**Figure 6g**). Quantification across TMA cores revealed a significantly higher proportion of Th17 (CD161⁺) T-cells among total CD3⁺ T-cells in AFR compared with EUR tumors specifically in Gleason 7 (4 + 3) samples (p = 0.0176), while the overall Th17 proportions decreased from Gleason 6 to Gleason 7 tumors (**Figure 6h**). Overall, these results demonstrate that Th17-mediated inflammation remains elevated in AFR PCa patients with intermediate-grade disease.

### Identification of Schwann Cells in the Prostate Tumor Microenvironment via Spatial and Single-Cell Transcriptomics

The large number of cells in our single cell and spatial datasets provided the ability to detect and characterize rare and previously unappreciated cell types in prostate cancers. We performed spatial transcriptomic profiling on two PCa samples, one AFR intermediate-risk localized PCa patient and one EUR localized PCa patient with perineural invasion (**Figure 7a,b**, **Supp Figure 8a).** After cell annotation, we identified a cluster of 2,526 cells in one sample with high expression of MPZ, SOX10, PMP22, and SCN7A consistent with Schwann cells (**Supp Figure 8b,c**). To validate the identity of these cells, we computed the signature score using a gene set of canonical Schwann cell markers and observed significant enrichment in the annotated Schwann cells in both samples (**Figure 7c**). The Schwann cells localized around nerve structures in the intermediate-risk localized PCa sample (**Figure 7d**, **Supp Figure 8d**), and in the perineural invasion sample, the Schwann cells were often surrounded by GLUT1⁺/NGFR⁺/MFAP5⁺ cells which are features consistent with satellite glial cells (**Supp Figure 8e**). The expression of GLUT1 in these perineural invasive areas suggests a change in metabolic activity, potentially driven by tumor-induced stress or hypoxia. Gene set enrichment analysis revealed that Schwann cells were enriched in pathways including glial cell differentiation, myelination, and Schwann cell development (**Supp Figure 8f**) and an enrichment of myogenesis signature, consistent with a mature and specialized glial phenotype. Our integrated single-cell RNA-seq atlas also contained a small cluster of 512 Schwann cells with up-regulation of neuronal markers MPZ, SCN7A, and ASPA compared with other cell types (**Figure 7e,f**). Each sample had less than 10 Schwann cells which were previously grouped with cells from the stromal compartment in a prior publication^70^. Our results suggest that spatial data and large-scale atlases can uncover rare cell types.

**Figure 8.**
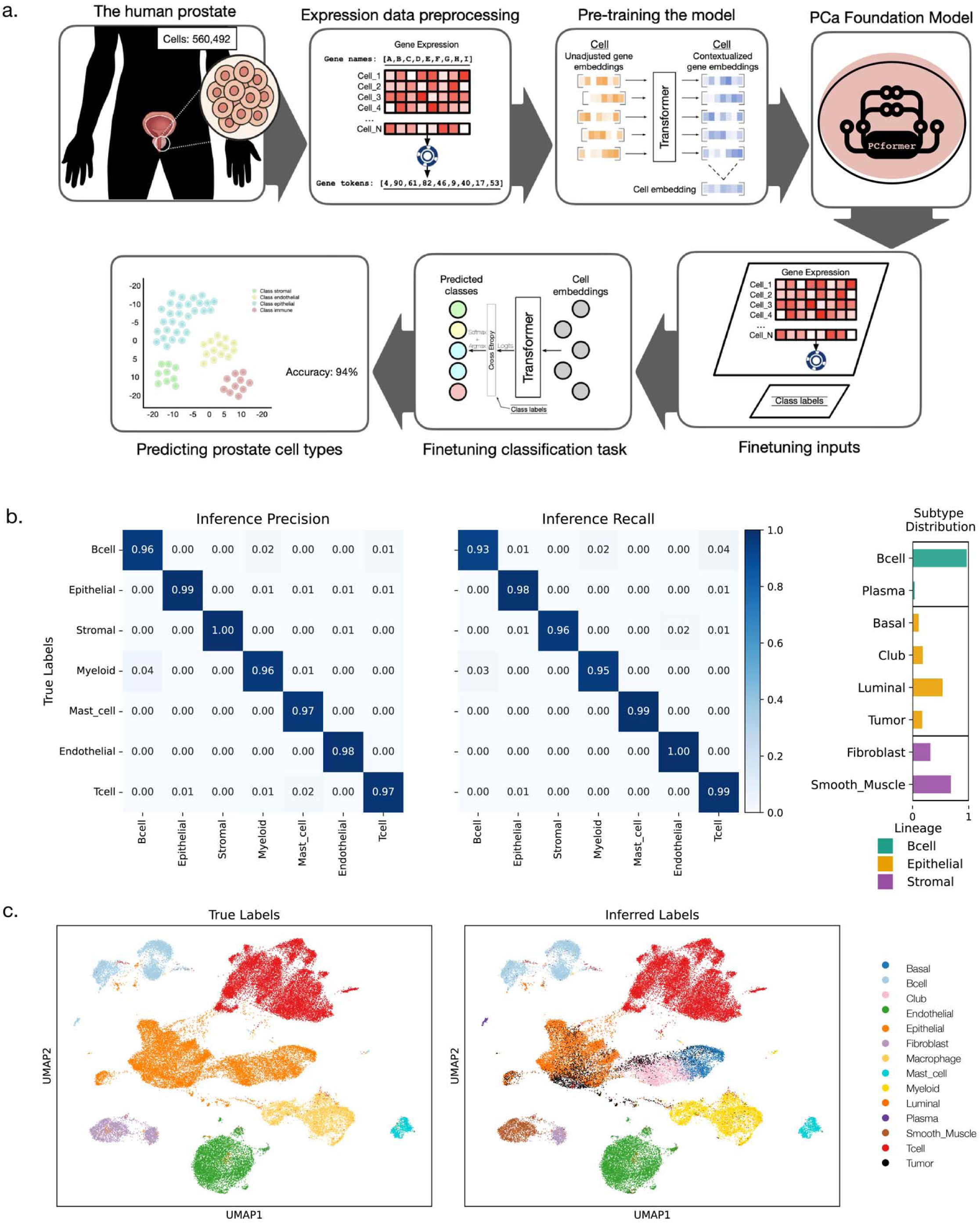
A high-performance transformer DL model for PCa. **a.** Schematic of PCformer workflow from data preprocessing to prediction. The PCformer transformer model is trained on ranked gene expression values from prostate single-cell RNA-seq data. Each input cell is represented as a sequence of gene tokens ordered by descending expression rank, with a maximum of 512 genes per cell. Tokens are embedded and combined with positional encodings before being passed through a stack of transformer encoder layers. During pretraining, PCformer learns to predict masked gene tokens using a self-supervised objective applied to over 500,000 cells from prostate tissues. This stage produces a generalized foundation model with contextual embeddings of prostate gene expression patterns. The foundation model is extended with a classification head and fine-tuned using cross-entropy loss on labeled datasets. During this step, the model adapts its weights to perform supervised prediction of prostate cell types. **b.** Confusion matrix showing the precision and recall results of inference on the Wong et al. dataset for each cell type. The published dataset did not contain annotations for smooth muscle, plasma, and epithelial subtypes. For visualization, these classes were collapsed into their respective lineages: Epithelial (basal, club, non-tumor luminal, and tumor), Stromal (fibroblast and smooth muscle), and B-cell (B-cell and plasma). The right panel shows the within lineage distribution of each subtype. **c.** UMAP showing the predicted cell type (right) and ground truth cell type labels (left) on the Wong et al. scRNA-seq dataset.

In our scRNA-seq PCa atlas, we then performed CellChat ligand-receptor analysis^71^ and found that Schwann cells showed ligand–receptor interactions with basal, club, tumor, and non-tumor epithelial populations (**Figure 7g**), including LAMB3–ITGA6 and LAMC2–ITGA6 interactions that have been previously associated with basement membrane remodeling^72–74^. Additionally, we observed interactions from epithelial cells to Schwann cells via GDF15–TGFBR2, indicating a potential role for TGF-beta signaling in regulating epithelial–nerve communication. Given recent studies implicating Schwann cell–epithelial crosstalk in TGF-beta signaling and nerve-associated tumor progression^75,76^, we next sought to evaluate this more directly using spatial transcriptomic data. Schwann cells showed high expression of TGF-beta receptors (e.g., TGFBR1, TGFBR2), while epithelial populations and tumor cells expressed the TGF-beta ligand GDF15 (**Supp Figure 8g**). These findings are consistent with our scRNA-seq-based predictions and support a Schwann cell–epithelial signaling axis mediated by the GDF15–TGFBR2 interaction and may have implications in mechanisms related to prostate cancer perineural invasion^75^.

### A high-performance transformer DL model for PCa

Given the heterogeneity among PCa tumor cells and the mix of methodologies used in previous scRNA-seq PCa studies, there is a need for a unified annotation method customized for PCa samples. Using our prostate single cell atlas data, we developed the Prostate Cancer Single-Cell Annotation Transformer (PCformer), a transformer model trained on prostate scRNA-seq datasets to predict tumor cells and the major TME cell types (**Supp Figure 9a**). The resulting foundation model was fine-tuned for a cell type classification task, allowing us to make predictions across 12 prostate cell type classes. To evaluate the model’s ability to generalize to previously unseen data, we evaluated its performance on a hold-out set of 56,050 cells, representing 10% of the PCa Atlas dataset (**Figure 8a**). The fine-tuned classifier achieved an overall accuracy of 0.94 across the 12 prostate cell types, with a macro F1-score of 0.94 indicating consistent performance across cell types regardless of class size (**Supp Figure 9b**). The model also maintained high average precision (0.94) and recall (0.94), showing a balanced trade-off between minimizing false positives and false negatives (**Supp Figure 9b**). The model showed strong classification performance on stromal and immune cell types, with F1 scores ranging from 0.93 to 1.00 (**Supp Figure 9b**). Tumor epithelial cells, non-tumor luminal, and club cells showed moderately lower F1-scores (0.87-0.88), potentially attributed to the mixed gene expression profiles from more continuous cell states^33,44^. Basal cells achieved an intermediate F1-score of 0.92. Receiver Operating Characteristic (ROC) analysis showed strong discriminative performance across all classes (mean AUC = 0.97; **Supp Figure 9c**). Together, these metrics demonstrate that PCformer maintains strong predictive accuracy across cell types even in the presence of class imbalance.

The model showed high classification performance on several cell types, particularly those with stromal and immune function (**Supp Figure 9b,c**). In the stromal group, predictions achieved strong metrics for endothelial cells (precision = 0.99, recall = 0.99, F1-score = 0.99), fibroblasts (precision = 0.98, recall = 0.96, F1-score = 0.97), smooth muscle (precision = 0.98, recall = 0.99, F1-score = 0.98). Predictions were slightly stronger (>=98%) for the immune cell types including mast cells (precision = 0.99, recall = 1.00, F1-score = 1.00), B-cells (precision = 1.00, recall = 0.99, F1-score = 1.00), T-cells (precision = 0.99, recall = 0.98, F1-score = 0.99), and myeloid cells (precision = 0.99, recall = 0.98, F1-score = 0.99). However, in that group the model showed higher difficulty predicting plasma cells (precision = 0.95, recall = 0.91, F1-score = 0.93). This cell type has a lower representation in the dataset (support = 298). The model showed moderately lower performance (F1-score 0.82-0.92) for the epithelial cells: tumor (precision = 0.87, recall = 0.86, F1-score = 0.87), non-tumor luminal (precision = 0.86, recall = 0.88, F1-score = 0.87), basal (precision = 0.91, recall = 0.93, F1-score = 0.92) and club cells (precision = 0.87, recall = 0.90, F1-score = 0.88) (**Supp Table 5**). This may be attributed to their transitional states and cellular plasticity, which create mixed gene expression profiles. Both basal and club cells are known to transition between different epithelial states, leading to intermediate phenotypes and overlapping transcriptional signatures^33,44^.

To evaluate the generalizability of PCformer, we performed validation using two independent scRNA-seq datasets^77,78^. The first dataset contained author-defined cell type annotations at a lower resolution than our model’s fine-grained labels. The reference grouped epithelial subtypes (basal, club, luminal, tumor) under a single epithelial category, fibroblasts with smooth muscle, and plasma cells with B-cells. For direct comparison, we collapsed model predictions to match the reference annotation hierarchy (**Figure 8b, Supp Figure9d**). The model achieved high performance across all collapsed cell types (precision > 0.96, recall > 0.93; **Figure 8b**) (**Supp Table 5**). A UMAP analysis shows cells predicted under the same label align together in the same cluster (**Figure 8c**). The model performance on this author-defined dataset was similar to the performance on our Atlas hold-out dataset (Supp **Figure 9e,f**). The second inference set containing 17,832 cells from the immune TME was annotated using the model predictions. Performance was the highest for endothelial, B cells, T cells, and mast cells (mean confidence > 0.98; **Supp Figure 9g,h**), similar to prediction confidence for these cell types in the Atlas hold-out inference (**Supp Figure 9f**). The model had high performance on other cell types, including club cells (mean 0.928), myeloid, plasma, basal (mean > 0.85), luminal (mean 0.817), and tumor (mean 0.792). Fibroblast cells and smooth muscle cells were predicted with lower confidence scores (mean 0.681 and 0.430, respectively). Ground-truth annotations were not available for this dataset; therefore, predictive accuracy could not be assessed. However, the confidence patterns were consistent with earlier findings and further support the model’s strong performance on independent datasets.

## Discussion

PCa is marked by profound molecular and cellular heterogeneity, which drives variability in therapeutic responses and clinical outcomes. In this work, we integrated scRNA-seq data from 128 patients to construct a harmonized atlas and elucidate the cell states and gene modules that contribute to heterogeneity in localized, castration-resistant, and metastatic disease.

We first characterized broad tumor cell states, including basal-like and club-like tumor cells. Basal-like tumor cells expressed markers such as TP63, which is associated with progenitor-like properties and increased resistance to androgen deprivation therapy (ADT)^33,44^. Basal-like tumor populations have been previously implicated in aggressive PCa subtypes and therapeutic resistance^20^, suggesting that these cells may play a role in treatment failure and disease progression. We also identified a novel population of club-like tumor cells characterized by high expression of LTF and LCN2. Although club-like tumor cells have not previously been described, they may be similar to other rare luminal progenitor cells that have been reported in the prostate epithelium^33^. Our large-scale atlas demonstrated that basal-like and club-like tumor cells are generally detected in lower proportions alongside of traditional luminal-like cells in localized PCa and were enriched for distinct transcriptional pathways compared to normal basal and club cells. One limitation is that our integration approach relied on the intersection of genes across 14 datasets, which resulted in the exclusion of some canonical markers. Another potential challenge is that the initial identification of cancer cells with copy number analysis can be difficult in cancer types with lower levels of aneuploidy such as PCa. By utilizing an additional dataset of normal luminal cells as a normal reference and combining inferCNV-based CNV scoring with a novel machine learning approach, our approach likely improved the accuracy for defining prostate cancer cells in these datasets and resolved inconsistencies among prior analyses^20,27^.

Each broad tumor cell state and individual cancer cell could be defined by combinations of gene co-expression modules. The luminal-like tumor cells expressed distinct combinations of luminal modules, each containing a different luminal marker, including *KLK2/3, KLK4/SPDEF, NKX3-1/DPP4, TMPRSS2/FOXA1, FOLH1/SPON2, MSMB, PSCA*, and *AR*, highlighting the existence of multiple luminal subtypes, on the contrary of previous assumption that luminal tumors are functionally uniform^29^. This heterogeneity suggests that while tumors may appear broadly luminal in morphology, their transcriptional properties and clinical outcome could differ significantly. In localized PCa for example, we observed a significant decrease in the number of luminal-associated modules in high-grade and high-stage disease. However, this did not occur as a uniform switch from a luminal-positive to a luminal-negative state. While aberrant lineage plasticity and infidelity have long been understood as a continues process^35^, we did not observe a standard progression within and across patients. Instead, individual tumor cells retained distinct combinations of luminal modules while losing others. This cell-to-cell variability produced a continuum of luminal integrity in cells across tumors with greater loss associated with more aggressive disease.

GPAS analysis identified gene modules with significantly elevated expression in metastatic disease including those enriched in specific organs as well as those shared across multiple sites^22^. Cell cycle modules were broadly elevated across many organ sites reflecting both the increased number of actively cycling cells within metastatic samples as well as the higher expression of these genes within the cells themselves. We also identified site-specific modules that contained transcriptional pathways more typical of primary tumors from that organ including osteoclast regulators in bone, neuro-migratory genes in brain, and erythroid-like programs in liver. These combinations of gene programs activated in metastatic cells reflects an increase of tumor-intrinsic growth capacities as well as adaptation with specific microenvironmental contexts. Certain metastatic sites, such as the brain and lymph nodes, had limited numbers of samples which potentially reduced the statistical power to find additional modules associated with these specific sites. Overall, our framework revealed a potential mechanistic basis for inter-lesion heterogeneity and with metastatic gene expression being the joint outcome of intrinsic growth-promoting programs and organ-specific selective pressures.

We further delineated the polylithic landscape of gene modules associated with NEPC. Specifically, we defined two “core” modules that were strongly associated with NEPC including a novel HES6 module as well as a module containing canonical NE markers. Additional “supporting” NE-associated modules related to cell proliferation and cytoskeleton remodeling were also identified and may help promote NE transitions. Importantly, individual cells from NEPC patients had distinct combinations of the core and supporting NE modules rather than adopting a linear path of differentiation where all NE-related genes were concordantly increasing along a single trajectory. To summarize the overall accumulation of the NE phenotype in individual cells, we developed a novel scoring system which discretized each NE-associated module into “low”, “intermediate”, or “high” expression, assigned a numerical score to each level, and summed the module scores together across all NE modules. This scoring system not only reflected the high levels of NE-modules in NEPC-annotated tumors but also revealed intermediate levels in subsets of cells from other tumors suggesting that NE transitions can begin at the molecular level before becoming histologically apparent.

To our knowledge we also generated the first ancestry-based analysis of prostate cancer single cell data to reveal differences in cell populations between EUR and AFR PCa patients. AFR patients exhibiting enrichment of heat shock protein (HSP)-expressing T-cells, potentially reflecting cellular stress from an unknown etiology. A potential caveat is all six AFR patients were sequenced on the Seq-well platform and the observed HSP upregulation in T-cells could be a result of technical factors associated with this platform. Future studies with independent AFR patient cohorts will be necessary to validate these finding and assess their clinical significance. AFR patients also exhibited an enrichment of Th17 pro-inflammatory T-cells, a population known for its role in immune modulation and tumor progression. These results were validated in an independent cohort of localized PCa where we observed the strongest differences between populations in men with intermediate risk tumors. Overall, these findings further support differences in immune microenvironmental dynamics in PCa from AFR men^79^.

Lastly, while other deep learning transformer models have been developed for single cell data^80–83^, we developed a model trained and fine-tuned on over 560,000 single cells from prostate cancers specifically, demonstrating the model’s performance for automated PCa scRNA-seq annotation and other potential future downstream tasks. In spite of the overall strong performance of PCformer, the model had somewhat lower performance for specific cell types. For example, the misclassification between tumor and non-tumor cells could be attributed to their shared lineage origins, as prostate tumors often arise from luminal progenitor cells with overlapping transcriptional profiles^84^. Additionally, Heterogeneity within prostate tumors could also contribute to misclassification as the continuous and combinatorial nature of plasticity and transdifferentiation could further blur the boundaries between categorial, discrete cell labels^84^. In summary, we established a comprehensive harmonized single-cell atlas of PCa for the study of tumor cell heterogeneity, patient classification, metastatic adaptation, and TME interactions. These findings advance our understanding of PCa at the single-cell level and offer a foundational resource for future studies on tumor evolution, therapy resistance, and biomarker discovery.

## STAR⍰METHODS

### KEY RESOURCES TABLE

#### Antibodies

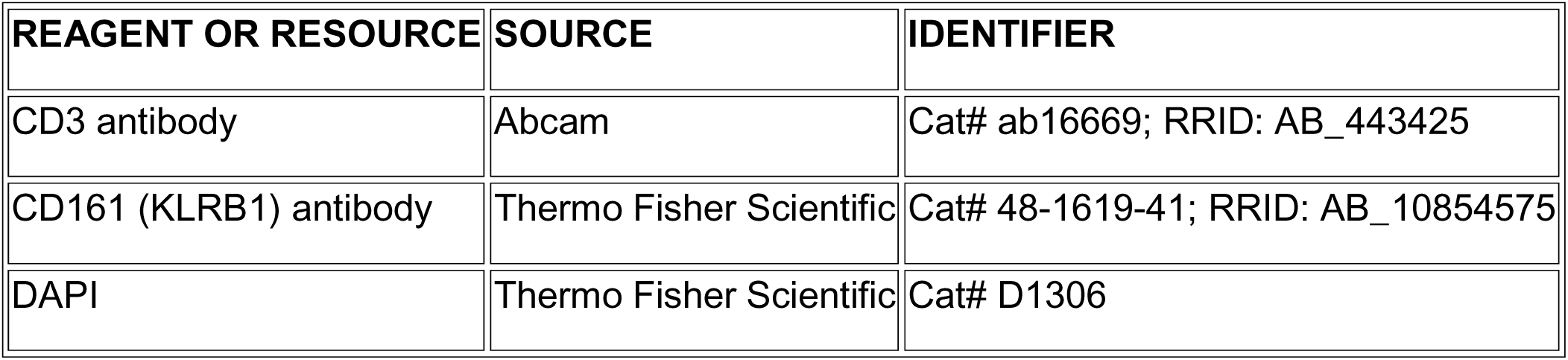

| REAGENT OR RESOURCE | SOURCE | IDENTIFIER |
| --- | --- | --- |
| CD3 antibody | Abcam | Cat# ab16669; RRID: AB_443425 |
| CD161 (KLRB1) antibody | Thermo Fisher Scientific | Cat# 48-1619-41; RRID: AB_10854575 |
| DAPI | Thermo Fisher Scientific | Cat# D1306 |

#### Biological samples

| REAGENT OR RESOURCE | SOURCE | IDENTIFIER |
| --- | --- | --- |
| Human prostate cancer tissue microarrays (TMAs) | Boston University School of Medicine | IRB-approved |
| Human metastatic prostate cancer FFPE tissue sections | UCSF and San Francisco Veterans Affairs Medical Center | IRB-approved |
| Human localized prostate cancer tissue (biopsies and RP specimens) | UCSF and San Francisco Veterans Affairs Medical Center | IRB-approved |

#### Chemicals, peptides, and recombinant proteins

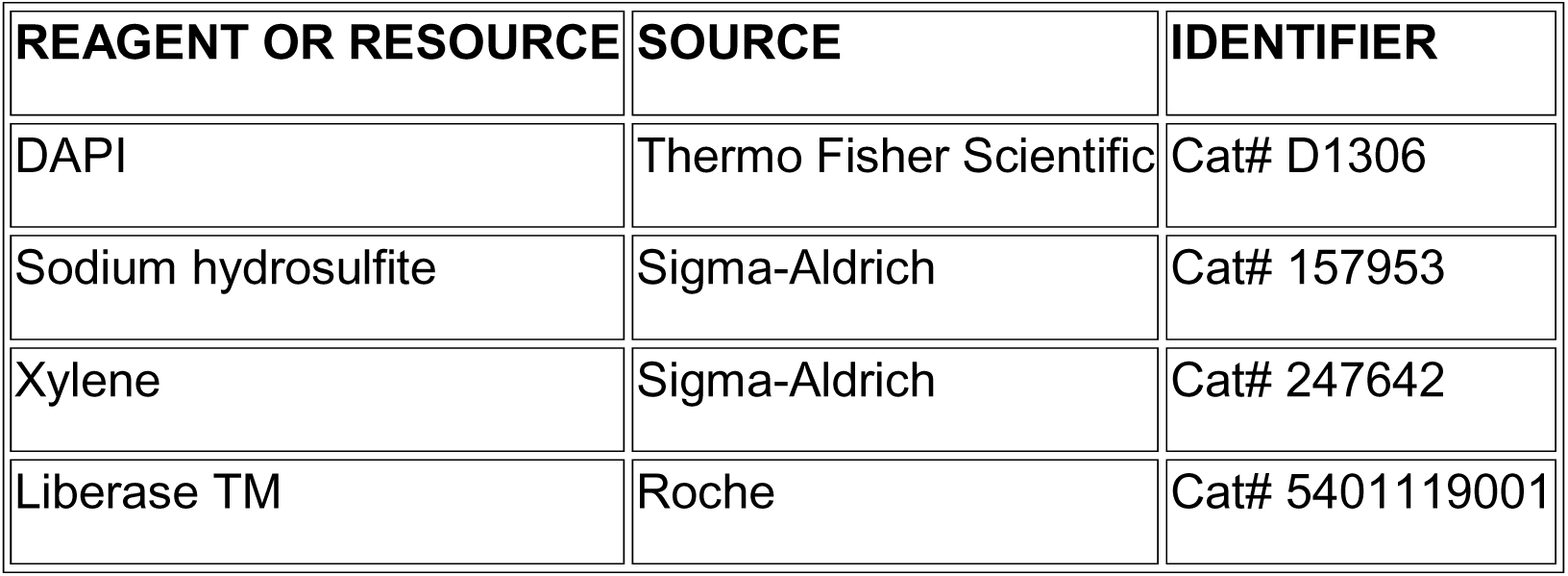

#### Commercial assays

| REAGENT OR RESOURCE | SOURCE | IDENTIFIER |
| --- | --- | --- |
| Seq-Well S3 barcoded mRNA capture beads | ChemGenes | MACOSKO-2011-10(V+) |
| Maxima H Minus Reverse Transcriptase | Thermo Fisher Scientific | EP0753 |
| KAPA HiFi HotStart ReadyMix PCR Kit | Kapa Biosystems | KK2602 |
| Nextera XT DNA Library Preparation Kit | Illumina | FC-131-1024 |
| 10x Genomics Chromium Single Cell 3' Kit | 10x Genomics | PN-1000691 |
| Xenium Prime 5K Human Pan Tissue & Pathways Panel | 10x Genomics | CG000578 |
| Xenium In Situ-Tissue Preparation Kit | 10x Genomics | CG000578 |
| Xenium Analyzer system | 10x Genomics | N/A |

#### Software and algorithms

| REAGENT OR RESOURCE | SOURCE | IDENTIFIER |
| --- | --- | --- |
| Seurat v5.2.1 | Stuart Lab | https://satijalab.org/seurat |
| Scanpy | Wolf et al. | https://scanpy.readthedocs.io |
| inferCNV | Broad Institute | https://github.com/broadinstitute/inferCNV |
| decontX | Celda package | https://bioconductor.org/packages/celda |
| celda | Celda package | https://bioconductor.org/packages/celda |
| QuPath | Bankhead et al. | https://qupath.github.io |
| XGBoost | Chen & Guestrin | https://xgboost.ai |
| PyTorch | Meta AI | https://pytorch.org |
| Xenium Explorer v3.2 | 10x Genomics | Proprietary |
| CUDA v12.6 | NVIDIA | https://developer.nvidia.com/cuda |

### EXPERIMENTAL MODEL AND STUDY PARTICIPANT DETAILS

#### Human prostate cancer samples

Human prostate cancer samples were obtained under Institutional Review Board–approved protocols at participating institutions, with informed consent obtained from all subjects. All samples were derived from male patients. Additional clinical and demographic characteristics are described in the main text and supplementary tables.

### METHOD DETAILS

#### Single cell data generation

Single-cell RNA sequencing was performed on a total of 18 prostate cancer samples. Sixteen samples — comprising six prostate biopsies and ten radical prostatectomy (RP) specimens from a prior cohort (patients 1–11) together with five additional RP localized prostate cancer samples — were processed using the Seq-Well S3 protocol^85,86^. Briefly, tissue samples were dissociated enzymatically using Liberase TM or collagenase type IV, filtered through 70-µm filters, and 10,000–20,000 cells were loaded onto arrays pre-loaded with ∼110,000 barcoded mRNA capture. Arrays were sealed with functionalized polycarbonate membranes and incubated at 37°C for 40 min, followed by lysis, hybridization, and bead collection as previously described. Reverse transcription was performed using Maxima H Minus Reverse Transcriptase overnight at 52°C, and whole transcriptome amplification (WTA) was carried out using the KAPA HiFi Hotstart Readymix PCR Kit. Libraries were prepared using the Nextera XT DNA Library Preparation Kit, dual-indexed, and sequenced on a NovaSeq S4 flow cell. Sequenced data were preprocessed and aligned using the dropseq_workflow on Terra (app.terra.bio), and digital gene expression matrices were generated for downstream analysis. The remaining two samples — one liver metastatic and one sphenoid metastatic prostate cancer sample — were processed using the 10x Genomics Chromium platform according to the manufacturer’s standard protocol and aligned using Cell Ranger v7.0^87^.

#### Data processing and integration

We compiled a comprehensive prostate cancer scRNA-seq atlas by combining samples from 12 published studies^18,20–22,24,27–33^ along with 9 unpublished samples. Each dataset underwent preprocessing, including quality control (QC) to filter out low-quality cells and genes. For published datasets, QC had been performed by the original authors; to ensure consistency across all datasets, we additionally applied a uniform filtering threshold, retaining only cells with 300 < nFeature_RNA < 10,000, nCount_RNA > 500, percent mitochondrial reads < 20%, and percent ribosomal reads < 50%. DecontX was used to estimate and remove ambient RNA within each sample^88^. Gene names were standardized across objects (replacing "." with "-"), and only genes detected in all datasets were retained for downstream analysis, yielding an intersection of 13,745 commonly detected genes. Integration and clustering was performed using the Seurat package. Decontaminated counts were normalized using the LogNormalize function. Highly variable genes were selected using the "vst" method with the top 3,000 variable features via FindVariableFeatures function. Integration anchor features were identified using SelectIntegrationFeatures function, and gene expression was scaled using the ScaleData function across the selected integration features prior to integration.

Integration was performed using the Reciprocal Principal Component Analysis (RPCA) algorith such that direct comparison of cells across samples and datasets originating from different studies or sequencing platforms is enabled. Cell clusters were identified using the Louvain algorithm via the FindClusters function, with clustering resolution optimized using the clustree package^89^. Clusters were annotated based on expression of canonical markers and signature gene sets, including epithelial, immune, stromal, and endothelial cells, with further sub-annotation of immune subsets (e.g., T-cells, myeloid cells, B-cells).

#### Tumor cell identification and tumor cell state analysis

Tumor cells were identified in a study-specific manner. For metastatic castration-resistant prostate cancer (mCRPC) samples from non-prostate sites, cells were first filtered to remove immune and stromal populations based on canonical marker gene expression (e.g., *PTPRC*, *CD3D*, *CD68*, *SELE, VWF, ACTA2*, *DCN*). Among the remaining epithelial cell clusters, tumor cells were identified at the patient level by the expression of known prostate cancer tumor markers (e.g., *ERG*, *FOLH1, SPON2, AMACR*) or prostate lineage markers (e.g., *NKX3-1*, *KLK3, KRT8*). Clusters meeting these criteria were designated as mCRPC tumor clusters for downstream analysis. For prostate tissue samples, epithelial cells from each dataset were analyzed with inferCNV^90^. The non-tumor cells from the corresponding dataset and luminal epithelial cells from the Henry et al. dataset^44^ were used as the reference set of cells. CNV scores were calculated as the squared sum of normalized and scaled CNV predictions extracted from the inferCNV results. High-confidence tumor cells defined as those a median absolute deviation (MAD) greater than 2.5 folds of MAD^91^.

To generalize tumor cell identification, we trained a separate multi-class XGBoost boosted gradient tree model^92^ within each dataset to classify high-confidence tumor cells versus other epithelial cells. Each model was run for 1,000 rounds with early stopping. Cells were then predicted as malignant or not by each individual dataset model. Cells predicted by at least three models were included in the final tumor cell dataset. Tumor sub-clusters were identified through unsupervised clustering of the tumor cells after integration with RPCA. Differentially expressed genes (DEGs) for each sub-cluster were computed using the FindAllMarkers function with the MAST algorithm to account for within-cluster variation^93^.

Pseudotime analysis was conducted using scanpy^94,95^, with the trajectory starting point defined by the highest stemness score based on CytoTRACE^96^. PAGA graphs were generated to visualize lineage relationships, and gene set enrichment analysis (GSEA) was performed on DEGs to identify enriched pathways and signatures^97^.

#### Gene module identification and analysis

Co-expression gene modules and cell clusters were co-clustered using the celda_CG algorithm from the celda package^34^. Initial 150 gene modules and 70 cell clusters for co-clustering was determined by the perplexity plot generated from the celda package using plotGridSearchPerplexity(). An additional 16 modules were split using the splitModule function resulting in a total of 166 modules. Modules were then reordered using hierarchical clustering using the reorderCelda function. Functional enrichment was performed for each module using Enrichr. Modules were annotated using a combination of enriched functional categories, known marker genes, and manual curation of genes. For gene program association studies (GPAS), module counts were normalized, log2 transformed, and analyzed using linear mixed effect (LME) models as previously described^26^. Briefly, our model used metastatic site as fixed variables and study, patient, and celda cluster as fully nested random effects. Module scores were corrected by subtracting the study coefficient for that module in each cell. Coefficients of the fixed variables were used as the log2 fold changes for each module at each metastatic site. P-value for each site were adjusted using the Benjamini-Hochberg False Discovery Rate (FDR).

#### Identification of gene modules associated with genomic alterations

The Cancer Genome Atlas Prostate Adenocarcinoma (TCGA PRAD) dataset was used to elucidate combinations of genomic aberrations (mutations and CNVs) associated with gene module expression^17^. 485 samples had both CNV and RNA-seq data. GISTIC2 was run on the TCGA PRAD dataset to identify genes with CNV loss or amplification^3,98^. The resulting filtered mutations and CNVs data were converted into binarized features, together with gene module scores, and were then used as inputs for CaDrA^99^. Genomic alterations with a frequency between 3% and 80% were included as potential features. For CaDrA, candidate search the top 5 starting points were evaluated and a maximum size of the meta feature of 5 individual features was chosen. Meta features were scored using the “ks_pval” function and the top scoring meta feature of each module that had an enrichment score above 0.35 and a significant FDR-corrected p-value were considered significant.

##### Multiplex immunofluorescence staining for Th17 T Cells

Multiplex immunofluorescence (mIF) staining was performed to identify Th17 T-cells in prostate cancer tissues. Two prostate cancer tissue microarrays (TMAs), comprising a total of 56 patient cores (29 African American (AFR) and 27 European American (EA)), were stained with antibodies against CD3 (T cell marker) and a Th17 T-cell marker CD161 (KLRB1).

Whole-slide images were acquired at 20× magnification. Qualitative and quantitative image analyses were performed using QuPath^100^, with cell detection and phenotyping across entire TMA cores. Representative high-magnification views were generated from regions of interest in each core.

#### PCformer data preprocessing and pretraining

We developed PCFormer, a prostate cancer-specific encoder-only genomic language model pretrained using masked language modeling on the labeled Atlas dataset. Cell types included immune (T-cells, B-cells, myeloid, mast, plasma), stromal (fibroblast, endothelial, smooth muscle), and epithelial subtypes (basal, club, non-tumor, tumor). Following Geneformer^102^, genes within each cell were ranked by expression and tokenized into a vocabulary of 13,745 genes. Genes with zero expression were removed prior to ranking to retain biologically informative features. The dataset was split into training (80%; 448,392 cells), validation (10%; 56,050 cells), and hold-out (10%; 56,050 cells).

Three special tokens were used: <PAD>, for sequence padding to a fixed length of 512; <MASK>, used for masked language modeling (MLM) by applying masking to 15% of tokens (80% replaced with <MASK>, 10% with random genes, and 10% unchanged); and <UNK>, for genes not present in the vocabulary. Token embeddings were represented using an embedding matrix with vocabulary size 13,748 and embedding dimension 1024, with positional encodings added to preserve gene rank information. RMS normalization^103^ (ε = 1x10⁻) was applied to stabilize activation magnitudes and reduce the risk of exploding or vanishing activations during training.

The model consisted of six encoder layers, each containing of a Multi-Head Self-Attention (MHSA) mechanism^104^ with eight attention heads and a Feedforward Neural Network^105^ (FFN) with 4x expansion and a Sigmoid Linear Unit (SiLU) activation function^106^. Linear layers used PyTorch’s default Kaiming Uniform initialization. Training used a batch size of 32 and sequence length of 512 with attention masking for padding tokens. Dropout (0.1) was applied to embeddings and attention layers. Optimization was performed using AdamW^107,108^ (learning rate 1x10⁻□, β₁ = 0.9, β₂ = 0.98, weight decay = 0.01), with 10% warm-up followed by cosine decay (0.5 cycles). The model was pretrained for five epochs.

Preprocessing required ∼5 hours and training ∼6 hours on a single NVIDIA RTX 4090 (48 GB VRAM) with an AMD EPYC 7C13 CPU and 128 GB RAM using PyTorch and CUDA 13.0. The pretrained encoder captured gene co-expression patterns and functional similarities.

#### PCformer fine-tuning for cell type classification

For classification, 504,440 cells (excluding hold-out) were split into training (90%; 453,996 cells) and validation (10%; 50,444 cells). Pretrained weights initialized the model, while the classification head was randomly initialized. Inputs were padded to 512 tokens, and the model was trained to predict 12 prostate cell types.

Fine-tuning used the AdamW optimizer with a learning rate of 3x10⁻□, β₁ = 0.9, β₂ = 0.999, and weight decay of 0.01, with a constant learning rate schedule. Training ran for five epochs using cross-entropy loss with class weighting to address imbalance. Sequence embeddings were aggregated using adaptive average pooling to produce a fixed-length cell representation, which was passed to a linear classification head and softmax for prediction.

#### PCformer inference and validation

Evaluation was performed on a 10% hold-out set (56,050 cells) and two external scRNA-seq datasets^77,78^. Preprocessing matched training conditions. During inference, the model operated in evaluation mode with batched inputs (length 512). Cell representations were obtained via adaptive average pooling and classified using a linear head with temperature-scaled softmax (T = 1.0) and argmax selection.

For external validation, the Wong et al. dataset^77^ was filtered to overlapping cell types (54,676 cells), while the Adorno-Febles dataset^78^ (17,832 cells) was evaluated without filtering due to lack of annotations.

#### Generation of Xenium 5K Prime data

Xenium sections were prepared following the Xenium *In Situ Tissue Preparation Guide protocol* (10X Genomics, Demonstrated Protocol, CG000578-Rev C). FFPE samples are sectioned at 5 μm thickness and placed on a 10X Xenium slide, dried for 3 hours at 42°C, and kept overnight in a vacuum desiccator. Slides are then baked for 30 mins at 60°C and deparaffinized with Neo-clear, ethanol, and nuclease-free water. Decrosslinking buffer is prepared with tissue enhancer, urea (final concentration of 0.5M) and diluted perm enzyme B in 1x PBS, and added to Xenium slides in a thermocycler with the following parameters: Decrosslinking at 80°C for 30 mins and re-equilibrated at 22°C for 10 mins. After 3 washes with 0.5X Saline-Sodium Citrate + 0.05% Tween-20, slides undergo polishing (polishing buffer, nuclease-free water and polishing enzyme) in a thermocycler at 37°C for 1 hour. After 3 washes with 1X PBS-0.05%Tween 20 (PBS-T), slides undergo overnight probe hybridization with the Xenium Prime 5K Human Pan Tissue & Pathways Panel in a thermocycler at 50°C for 18 hours. After overnight incubation and per protocol washes with PBS-T, priming hybridization mix is applied to the slides and incubated in a thermocycler at 50°C for 1:30 hours for priming and 30 mins for post-priming wash with the post-priming hybridization wash buffer. Slides are treated with RNase at 37°C for 20 mins in a thermocycler. After appropriate per protocol washes, post-hybridization wash is added to slides and incubated for 15 mins at 35°C in a thermocycler followed by ligation at 42°C for 30 mins in the thermocycler. After appropriate per protocol washes, an amplification enhancer mix is applied to the slides and incubated for 2 hours at 4°C in a thermocycler followed by an amplification enhancer wash buffer. Amplification master mix is then applied to slides and incubated for 1.5 hours at 30°C. Cell boundary staining is next performed with the Xenium Block and Buffer stain and the Xenium Multi-Tissue stain mix and slides are incubated overnight for 18 hours at 4°C. Cell markers for cell boundary staining used are ATP1A1, CD45 and E-Cadherin. The following day, slides undergo auto-fluorescence quenching and nuclei staining before being loaded onto the 10X Xenium analyzer.

##### Xenium Analyzer

The Xenium Analyzer is a robust imaging based sequencing platform enabling high-throughput identification of RNA targets at sub-cellular resolution with XY localization. The Xenium Analyzer was run following the 10X Genomics Demonstrated Protocol CG000584. Decoding Module A and B were thawed at 4℃ overnight followed by a 300 RCF spin cycle prior to loading onto the Xenium Analyzer. Deionized water, Xenium Sample Wash Buffer A (Nuclease Free Water 895 mL; 10X PBS 100 mL; 10% Tween 20 5mL) and B (Milli-Q Water 500 mL), and Xenium Probe Removal Buffer (Nuclease Free Water 139.5 mL; DMSO 100% 150 mL for final concentration of 50%; KCL 2000 mM 7.5 mL for final concentration of 50 mM and Tween 20 10% 3mL for final concentration of 0.1%); are prepared and loaded onto the Xenium Analyzer. Slides were placed on the Xenium Analyzer and the user manually selected the tissue region on a low-resolution full slide image. Data collection occurs in different cycles of fluorescently labeled probe binding followed by image acquisition and stripping of probes. Samples had additional fluorescent images taken on the Xenium multi-tissue stain channel to better perform cell segmentation. Run time on the 10X Xenium analyzer was 4 days.

#### Hematoxylin and Eosin staining of Xenium slides

Slides were removed from the Xenium Analyzer and underwent quencher removal with sodium hydrosulfite (Sigma-Aldrich 157953) at a final concentration of 10mM according to 10X Genomics’ Demonstrated Protocol CG000613. Immediately after quencher removal, slides underwent conventional Hematoxylin and Eosin staining according to the following protocol: Distilled H20 rinse for 30 sec, Hematoxylin (Modified Mayer’s Hematoxylin StatLab/American Mastertech HXMMHGAL) rinse for 1 min, Hematoxylin incubation for 11 min and 30 sec, tap water rinse for 1 min, Bluing solution (Scott’s Tap Water with Magnesium Sulfate) rinse for 1 min, tap water rinse for 1 min, Ethanol 70% rinse for 1 min, Eosin Y with Phloxine B (StatLab/American Mastertech STE0257) for 30 sec, Ethanol 70% rinse for 15 sec, Ethanol 95% rinse for 15 sec, Ethanol 95% rinse for 15 sec twice, Ethanol 100% rinse for 1 min, Ethanol 100% rinse for 1 min and 30 sec and Xylene (Sigma-Aldrich 247642) rinse for 2 min twice. The H&E stainer was linked to a tissue-Tek Film coverslipper and coverslipping was performed at room temperature immediately following the staining (Tissue-Tek coverslipping film 4770). Xenium Explorer is an interactive desktop software allowing users to interactively visualize RNA transcripts in tissue at subcellular resolution. Post Xenium H&E image alignment with DAPI stained cell segmentation was performed on Xenium Explorer version 3.2.

#### Xenium 5K Prime data processing and analysis

Cells with less than 50 total counts were excluded. Additionally, cells with more than 2,000 counts or more than 1,000 gene detected were examined on the H&E stain using the Xenium Explorer and removed after being identified as doublets. 508,742 cells were subsequently clustered and annotated using Seurat (Version 5.2.1). Cell type annotation was achieved using the FindTransferAnchors and TransferData functions in Seurat with a previously annotated localized PCa single-cell RNA-seq data as reference. Annotatons were confirmed then using canonical markers and established signature gene sets. Once cell types were identified and annotated, we overlaid the spatial cell type distribution onto the stained H&E image with the Xenium Explorer 3.2 software. The basal-like and club-like tumor cells were validated using two samples include one previously described localized prostate cancer FFPE sample and one publicly available sample from the FFPE Human Prostate Adenocarcinoma with 5K Human Pan Tissue and Pathways Panel (https://www.10xgenomics.com/datasets/xenium-prime-ffpe-human-prostate). Tumor cells were annotated using the transfer label method. Basal-like, club-like and luminal tumor signature scores were computed via the AddModuleScore function using previously establish signature gene sets. Then basal-like and club-like tumor cells were identified as tumor cells with outlier high basal-like and club-like scores (mean + 2*standard deviation). Matching H&E images were also obtained, and screenshots were taken for zoom-in views of regions for these basal-like and club-like tumor cells. The tumor identities of these cells were later confirmed by pathologists at Boston University and University of California San Francisco.

### QUANTIFICATION AND STATISTICAL ANALYSIS

Statistical analyses and quantification methods are described within the corresponding Method Details sections and figure legends. Sample sizes (n), definitions of replicates, and statistical tests used are reported in the figures and Results. Multiple hypothesis testing was corrected using false discovery rate where applicable. No data were excluded unless failing predefined quality control criteria.

## ACKNOWLEDGMENT

This work was conducted as part of the National Cancer Institute (NCI) Center to Reduce Cancer Health Disparities (CRCHD) R01CA248920 to J.D.C. and F.W.H. This work was also supported by Biohub San Francisco, Benioff Initiative for Prostate Cancer Research, NIH (U54CA302452), Prostate Cancer Foundation Challenge Award to F.W.H.

## ADDITIONAL RESOURCES

This scRNA-seq PCa atlas dataset integrated 14 datasets. 3 of 14 datasets were generated in the Huang Lab and the other 11 were downloaded from previously published studied. Data_1 referred to the Dong et al. study (Single-cell analysis supports a luminal-neuroendocrine transdifferentiation in human prostate cancer) and the processed data was downloaded from https://www.ncbi.nlm.nih.gov/geo/query/acc.cgi?acc=GSE137829. Data_2 referred to the Cheng et al. study (Pre-existing Castration-resistant Prostate Cancer–like Cells in Primary Prostate Cancer Promote Resistance to Hormonal Therapy) and the individual sample-specific gene expression matrices were downloaded from https://www.ncbi.nlm.nih.gov/biosample?LinkName=bioproject_biosample_all&from_uid=699369 (only the human samples). Data_3 referred to the Ci et al. study (Single-Cell RNA-seq Reveals a Developmental Hierarchy Super-Imposed Over Subclonal Evolution in the Cellular Ecosystem of Prostate Cancer) and the processed .rda file was downloaded from https://ngdc.cncb.ac.cn/gsa-human/browse/HRA000823 with the consent from the corresponding author of this study. Data_4 referred to the Tuong et al. study (Resolving the immune landscape of human prostate at a single cell level in health and cancer) and the processed and annotated data object was downloaded from https://www.prostatecellatlas.org. Data_5 referred to the Heidegger et al. study (Comprehensive characterization of the prostate tumor microenvironment identifies CXCR4/CXCL12 crosstalk as a novel antiangiogenic therapeutic target in prostate cancer) and the gene expression matrices were downloaded from the GEO repository with the accession number GSE193337 (secure token: srgpkcwkvvejfgt). Data_6 referred to the Karthaus et al. study (Regenerative potential of prostate luminal cells revealed by single-cell analysis) and the gene expression matrix .h5 file was downloaded from https://www.ncbi.nlm.nih.gov/geo/query/acc.cgi?acc=GSE146811. Data_7 referred to the Chen et al. study (Single cell analysis reveals onset of multiple progression associated transcriptomic remodellings in prostate cancer) and the gene expression matrix was downloaded from https://www.ncbi.nlm.nih.gov/geo/query/acc.cgi?acc=GSE141445 and the annotation metadata was obtained from the original manuscript. Data_8, Data_9 and Data_10 referred to the three in-house datasets of 16 localized PCa samples and two mCRPC samples. Raw FASTQ files and gene expression matrices have been deposited in the GEO repository with the accession number GSE324131 (secure token: anahssocxzkxfwz). Data_11 referred to the Ma et al. study (Identification of a distinct luminal subgroup diagnosing and stratifying early stage prostate cancer by tissue-based single-cell RNA sequencing) and the gene expression matrix was downloaded from the GEO repository with the accession number GSE157703. Data_12 referred to the Chan et al. study (Lineage plasticity in prostate cancer depends on JAK/STAT inflammatory signaling) and the gene expression matrix was download from the GEO repository with the accession number GSE210358. Data_13 referred to the Hirz et al. study (Dissecting the immune suppressive human prostate tumor microenvironment via integrated single-cell and spatial transcriptomic analyses) and the gene expression matrix with annotation metadata were downloaded from the GEO repository with the accession number GSE181294. Data_14 referred to the Kfoury et al. study (Human prostate cancer bone metastases have an actionable immunosuppressive microenvironment) and the gene expression matrix was downloaded from the GEO repository with the accession number GSE143791. Two other datasets were also used in this analysis to test the accuracy of our transformer model including the Wong et al. dataset (Single cell analysis of cribriform prostate cancer reveals cell intrinsic and tumor microenvironmental pathways of aggressive disease) and the Victor et al. dataset (Single-cell analysis of localized prostate cancer patients links high Gleason score with an immunosuppressive profile). Gene expression matrices were downloaded for these two datasets and Wong et al. cell type annotation was acquired from https://github.com/shengqh/Hurley2022scRNA/tree/main.

