## Supplementary Figures for "A single-cell and spatial atlas of prostate cancer reveals the combinatorial nature of gene modules underlying lineage plasticity and metastasis"

### Supp Figures

Supp Figure 1

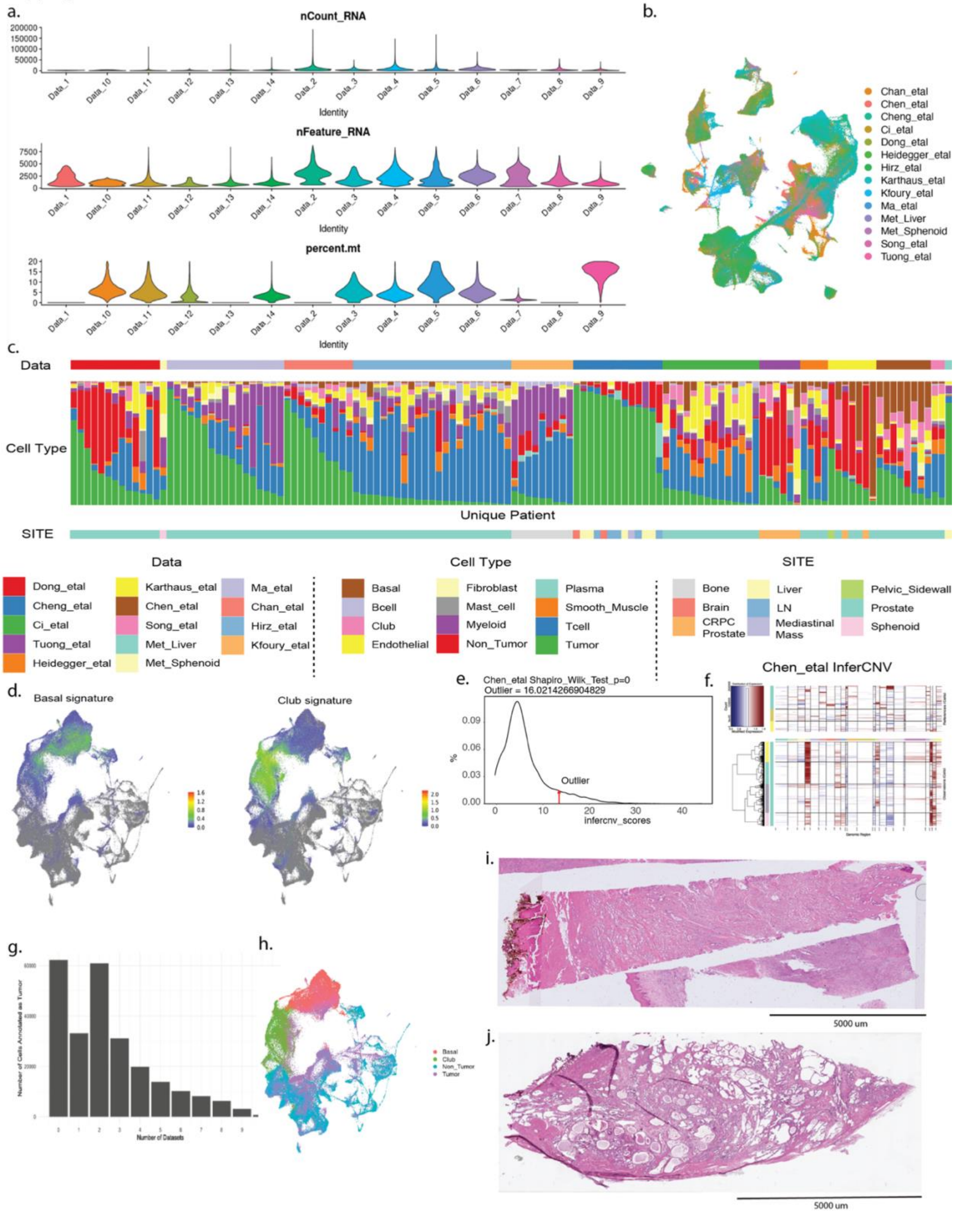

**Supp Figure 1. Quality control and integration metrics for the PCa scRNA-seq atlas**

**a.** Violin plots showing the distribution of nCount_RNA, nFeature_RNA, and percent mitochondrial reads across each dataset included in the integration. **b.** UMAP plot colored by dataset ID demonstrating integration of samples. **c.** Stacked bar plot showing cell type composition across unique patients, colored by cell type annotation with site annotation shown below. **d.** Feature plots showing basal and club epithelial signature scores projected onto the integrated UMAP. **e.** Distribution of copy number scores for all the epithelial cells and the red arrow marks the threshold above which cells are considered to harbor outlier high copy number changes. **f.** Example inferCNV heatmap for one dataset in the atlas using all the non-tumor epithelial cells as reference (up) and inferred copy number profiles for the unknown cells (down). Copy numbers are represented in a blue-red color scheme (blue: deletion, red: amplification). **g.** Bar plot showing the number of predicted tumor cells by different model training results. **h.** UMAP visualization of epithelial cells colored by epithelial subtype. **i**. H&E staining image of the first FFPE localized prostate cancer sample from a Gleason 3+3 localized prostate cancer for spatial profiling using the Xenium Prime 5k gene panel. **j**. H&E staining image of the second FFPE localized prostate cancer sample from a Gleason 3+3 localized prostate cancer for spatial profiling using the Xenium Prime 5k gene panel.

Supp Figure 2

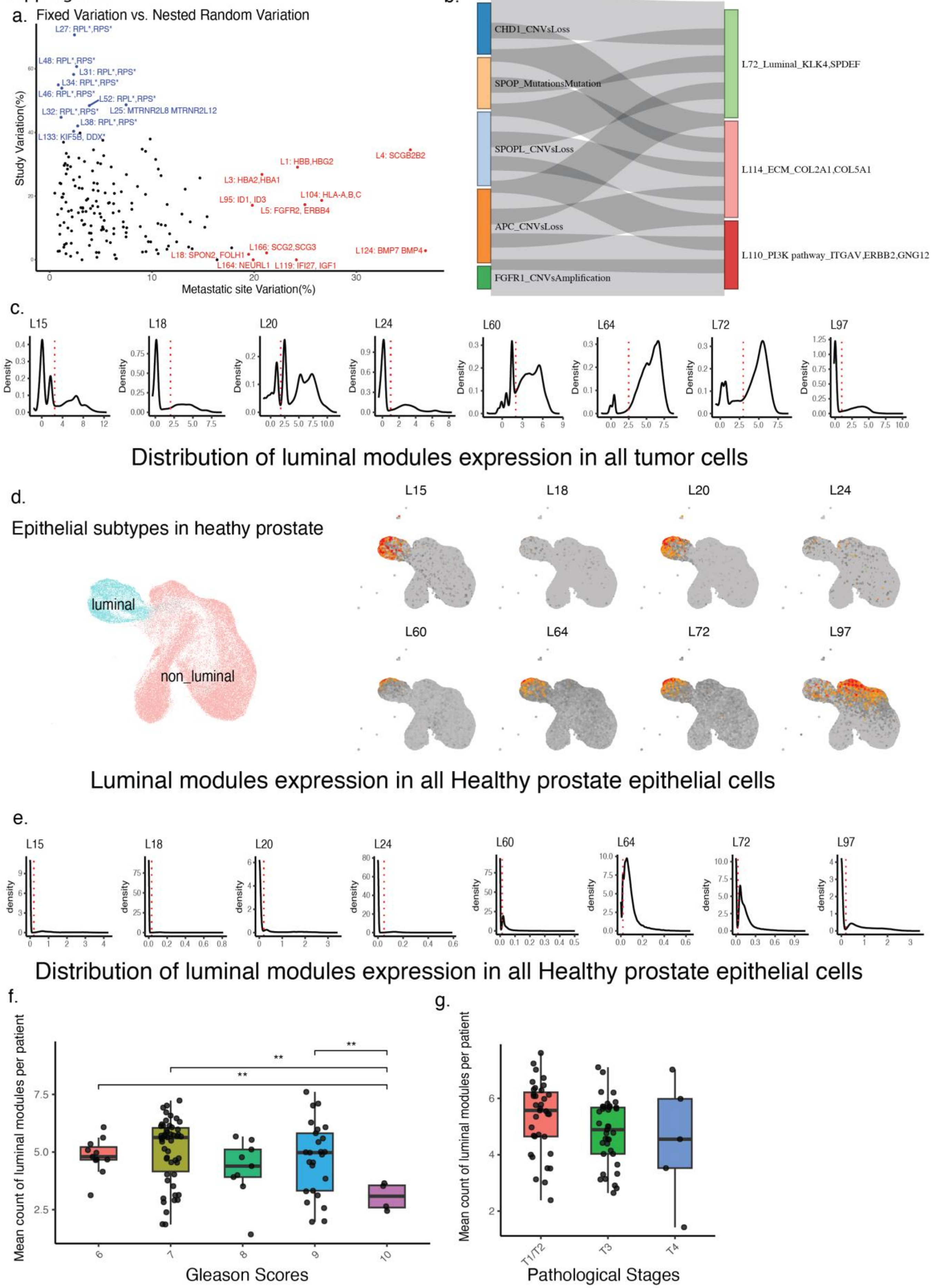

**Supp Figure 2. Classification of gene modules association and luminal module expression subtyping.**

**a.** Site variation percentage vs study variation percentage for each gene co-expression module. **b.** Connection between genetic mutation and different module expression in TCGA. **c.** Density plot of luminal modules expression in all tumor cells. **d.** Luminal modules expression in all Healthy prostate epithelial cells shown in UMAP. **e.** Density plot of luminal modules expression in all Healthy prostate epithelial cells. **f.** Association between Gleason score and average number of cellular expressed luminal modules per patient. **g.** Association between tumor stage and average number of cellular expressed luminal modules per patient.

Supp Figure 3

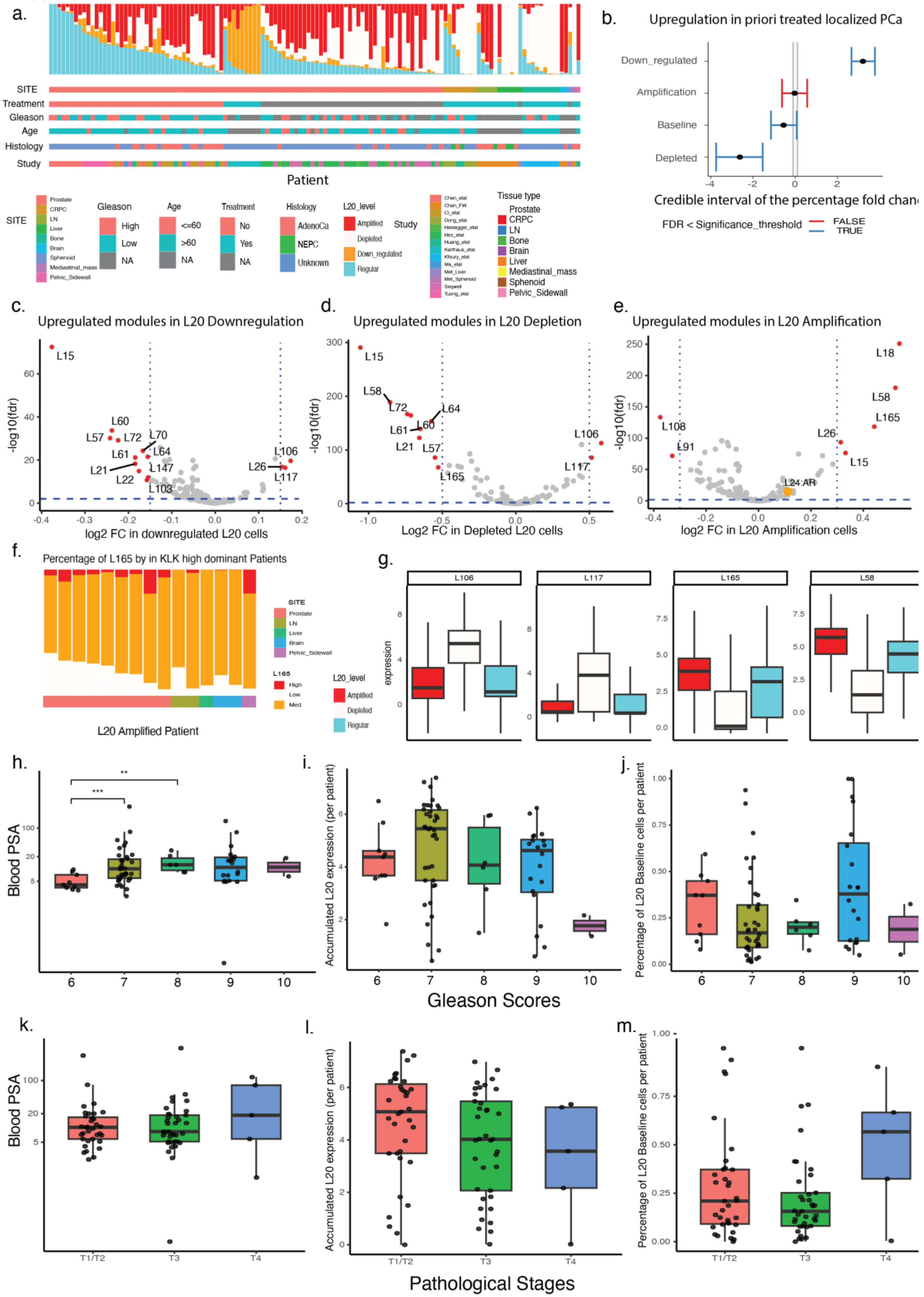

**Supp Figure 3. Delineating L20(KLK2/3) module expression categories and clinical association**

**a.** Spectrum of composition of L20(KLK2/3) expression categories across patient labelled by clinical metadata. **b.** Credible interval of the percentage fold change for different L20 expression cell types in different localized prostate cancer from sccomp analysis. **c-e** volcano plot of differential module expression between L20 down-regulation, L20 depleted and L20 amplification cells compared to L20 baseline. **f.** Composition of L165(MS4A8, HOXA10) expression subtypes in patients with >75% L20- amplification cells. **g.** Cellular module expression of L106, L117, L165 and L57 in different L20 expression subtypes. **h-j** Association between Gleason score and blood PSA, accumulated L20 expression and percent L20 baseline tumor cells. **k-m** Association between tumor stage and blood PSA, accumulated L20 expression and percent L20 baseline tumor cells.

Supp Figure 4

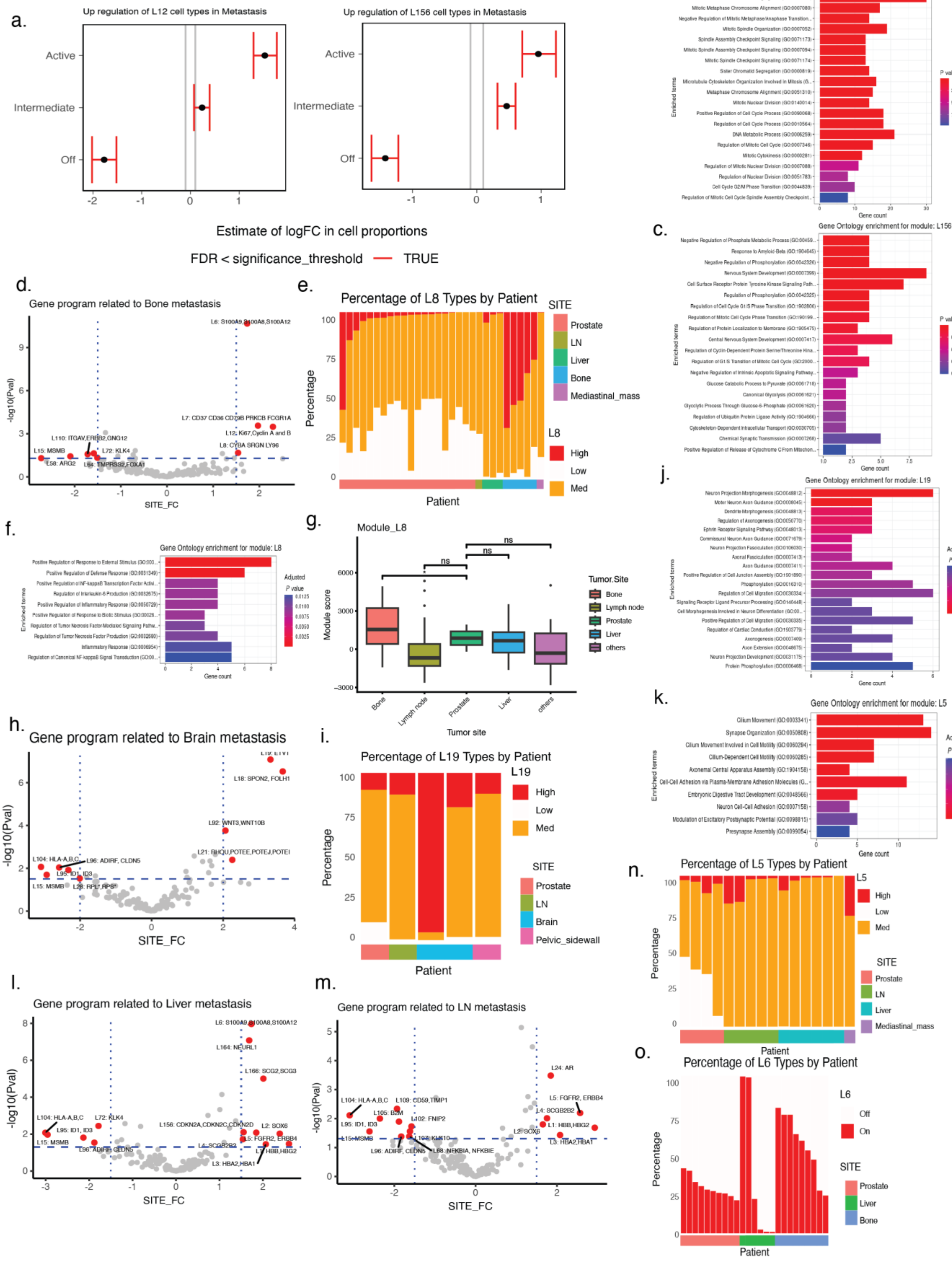

**Supp Figure 4. Dissecting biological functions of metastasis associated modules.**

**a.** Credible interval of the percentage fold change for different L12 and L156 expression cell types in metastasis vs localized prostate cancer from sccomp analysis. **b.** Gene Ontology enrichment for genes in module L12. **c.** Gene Ontology enrichment for genes in module L156. **d.** Volcano plot of differential module expression in liver metastasis compared to localized PCa. **e.** Composition of L6 expression cell types in different patient. **f.** Volcano plot of differential module expression in bone metastasis compared to localized PCa. **g.** Composition of L8 expression cell types in different patient. **h.** Gene Ontology enrichment for genes in module L8. **i.** Module Expression score of L8 in different metastatic sites in SU2C dataset. **j.** Volcano plot of differential module expression in Brain metastasis compared to localized PCa. **k.** Composition of L19 expression cell types in different patient. **l.** Gene Ontology enrichment for genes in module L19. **m.** Volcano plot of differential module expression in lymph nodes metastasis compared to localized PCa. **n.** Composition of L5 expression cell types in different patient. **o.** Gene Ontology enrichment for genes in module L5. **p.** Module Expression score of L8 in different metastatic sites in SU2C dataset.

Supp Figure 5

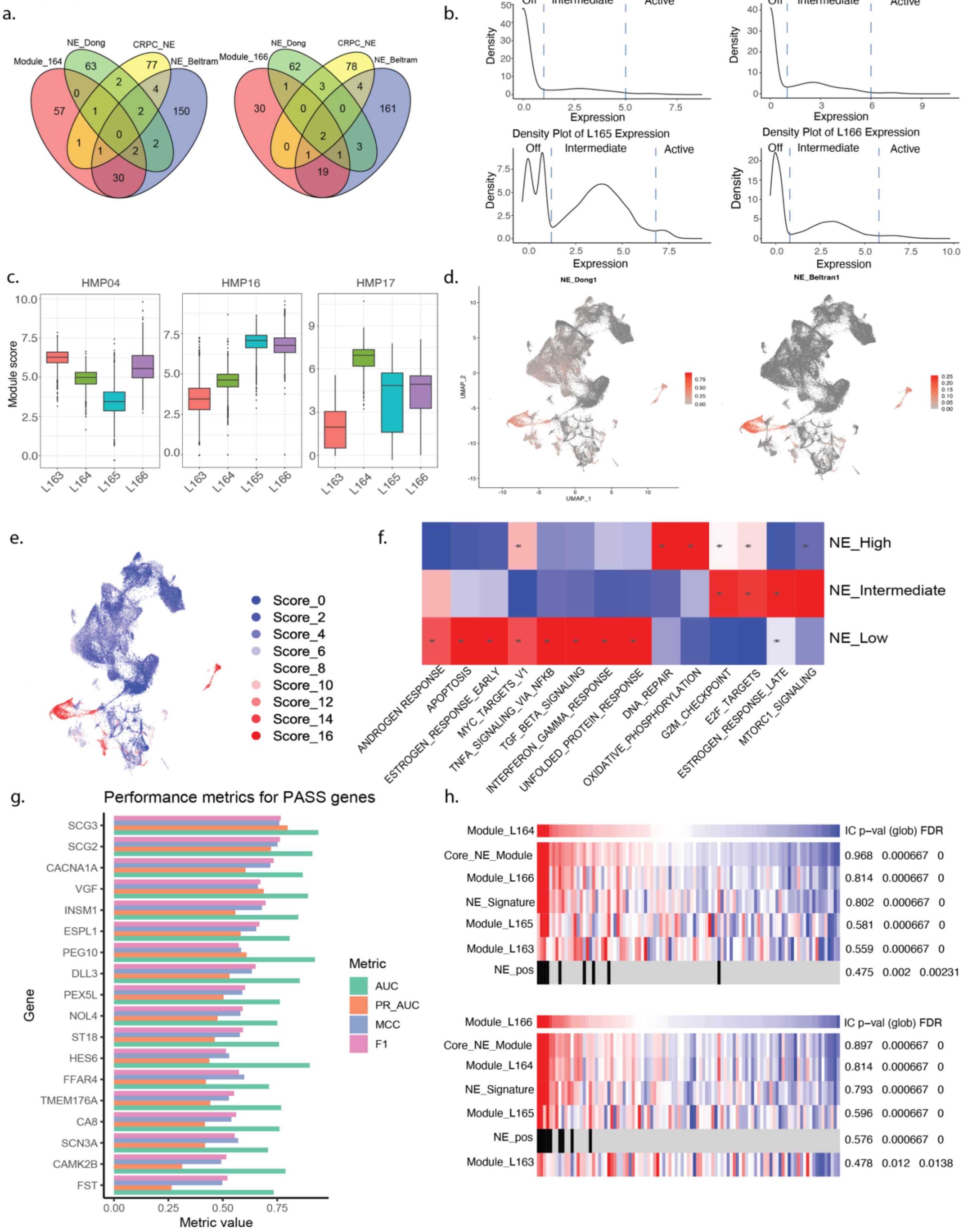

**Supp Figure 5. Neuroendocrine-like module analysis across PCa datasets**

**a.** Venn diagrams showing gene overlapping between Modules L164 and L166 and previously established NE signatures (NE_Dong, NE_Beltran, CRPC_NE). **b.** Density plot of module scores across tumor cells for Modules L163-L166, classified into three categories, Off (low), Intermediate, and Active (high) based on the distribution. **c.** Box plots showing module scores for L163-L166 across tumor cells from three NEPC patients (HMP04, HMP16, HMP17). **d.** Feature plots showing neuroendocrine signature scores (NE_Dong, NE_Beltran) across tumor cells, highlighting enrichment in subsets of NE-like cells. **e.** UMAP of tumor cells colored by NE scores (0–16), demonstrating a gradient distribution of NE module scores across the tumor cell population. **f.** Gene set enrichement analysis result showing the significantly enriched pathways for NE_low, NE_intermediate and NE_high gene sets. **g.** Bar chart summarizing performance metrics (AUC, PR-AUC, MCC, F1) for the 18-gene NE. h**.** ssGSEA heatmaps of NE-associated between the two core lineage NE modules (L164 and L164) and other NE signatures/modules in the SU2C dataset, showing information content (IC), association p-values, and FDR for each signature.

Supp Figure 6

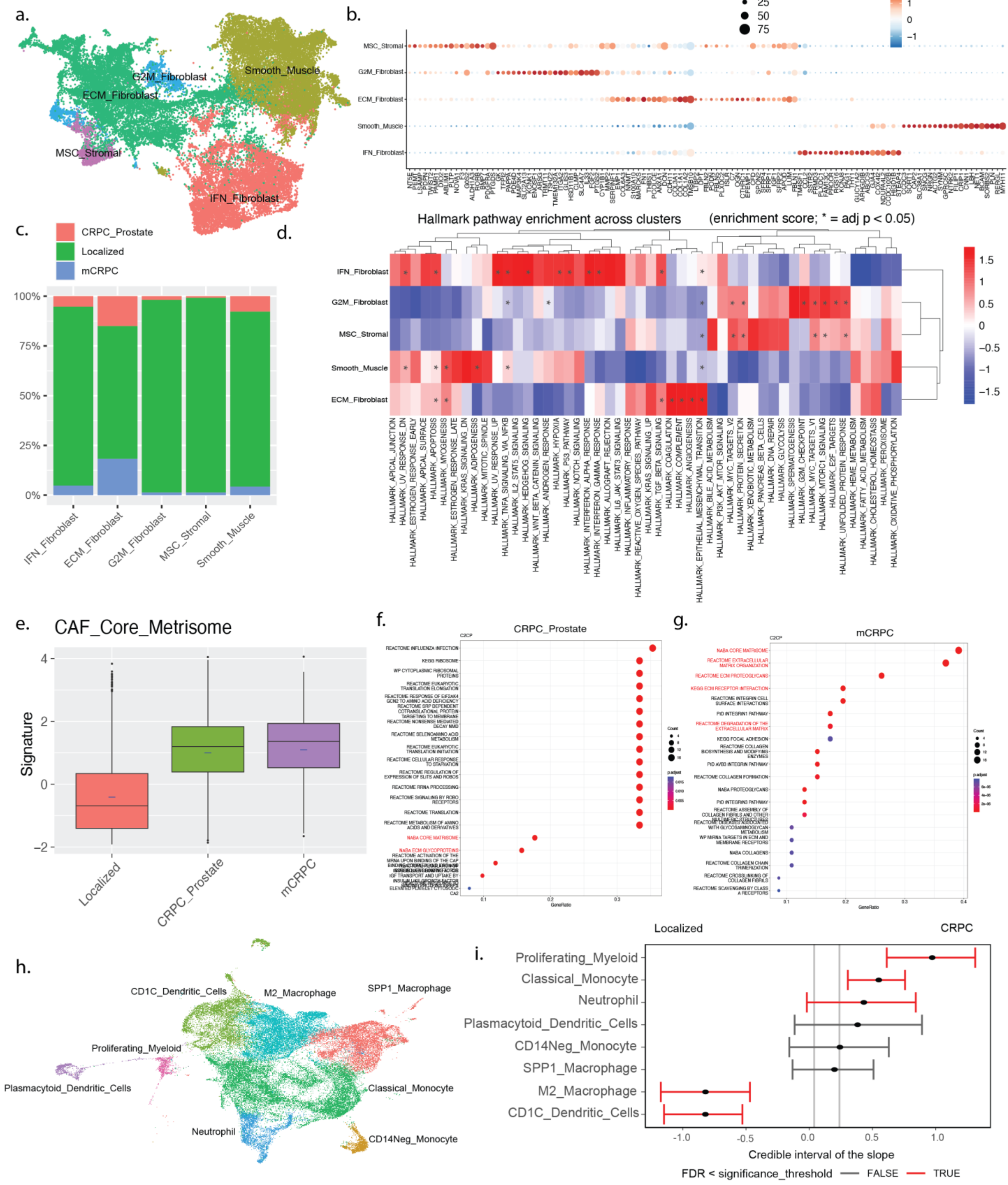

**Supp Figure 6. Clustering and characterization of stromal cell populations in the PCa scRNA-seq atlas**

**a.** UMAP plot showing stromal clustering results with clusters annotated as ECM fibroblasts, MSC stromal cells, smooth muscle cells, G2M fibroblasts, and IFN fibroblasts. **b.** Dot plot of selected marker genes across stromal clusters with dot size indicating percent expression and color indicating average expression. **c.** Stacked bar plot showing stromal cluster composition across localized, CRPC prostate, and mCRPC samples. **d.** Heatmap showing hallmark pathway enrichment across stromal clusters (enrichment score; * = adj p < 0.05). **e.** Boxplot showing CAF core matrisome signature scores across localized, CRPC prostate, and mCRPC stromal cells. **f-g.** Dot plots showing Reactome pathway enrichment analysis for differentially expressed genes in CRPC prostate (h) and mCRPC (i) compared to localized samples. **h.** UMAP plot showing clustering results for myeloid populations including CD1C_Dendritic_Cells, plasmacytoid dendritic cells, classical monocytes, CD14Neg monocytes, M2 macrophages, SPP1 macrophages, neutrophils, and proliferating myeloid cells. **i.** Compositional analysis of myeloid populations comparing localized prostate cancer and CRPC patients using sccomp. Error bars representing credible intervals; populations with FDR below the significance threshold are highlighted.

Supp Figure 7

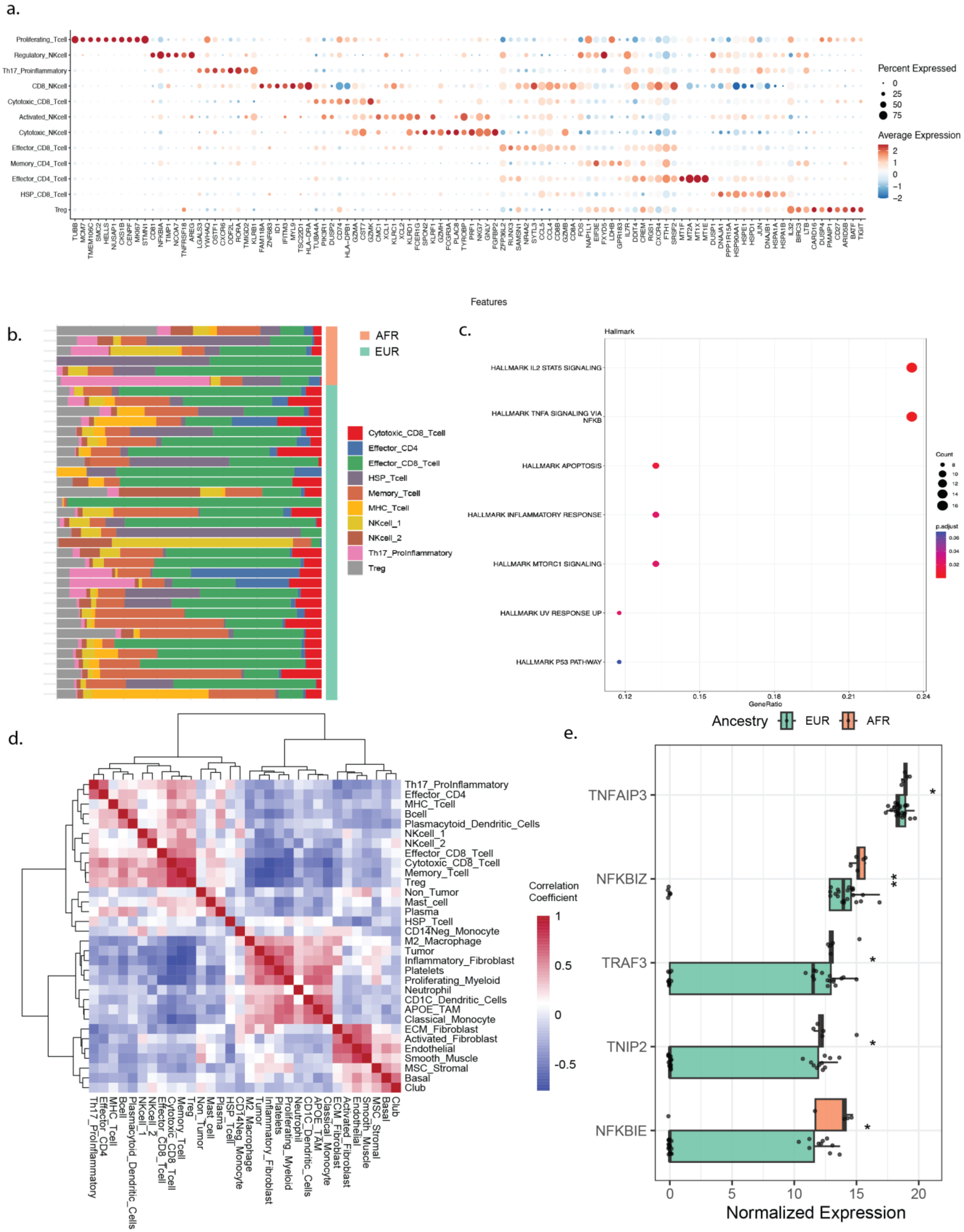

**Supp Figure 7. T-cell cell subpopulation signatures and analyses**

**a.** Dot plot showing expression of marker genes across T-cell subclusters. Dot size represents the percentage of cells expressing each gene, and color intensity indicates average expression. **b.** Patient-level T-cell subtype composition across AFR and EUR patients. Stacked bar plots illustrate relative proportions of T-cell subtypes for each patient. **c.** Gene set enrichment analysis of Th17 T-cell marker genes showing enrichment of IL2–STAT5 signaling, TNFα signaling via NFκB, inflammatory response, apoptosis, and related pathways. **d.** Correlation matrix showing relationships between immune and stromal cell populations across patients. Th17 T-cells positively correlate with multiple T-cell populations and are negatively correlated with stromal components including ECM and activated fibroblasts. **e.** Expression of NFκB-related genes (*TNFAIP3*, *NFKBIZ*, *TRAF3*, *TNIP2*, *NFKBIE*) in Th17 T cells from AFR versus EUR patients, supporting activation of TNFα–NFκB signaling in AFR tumors.

Supp Figure 8

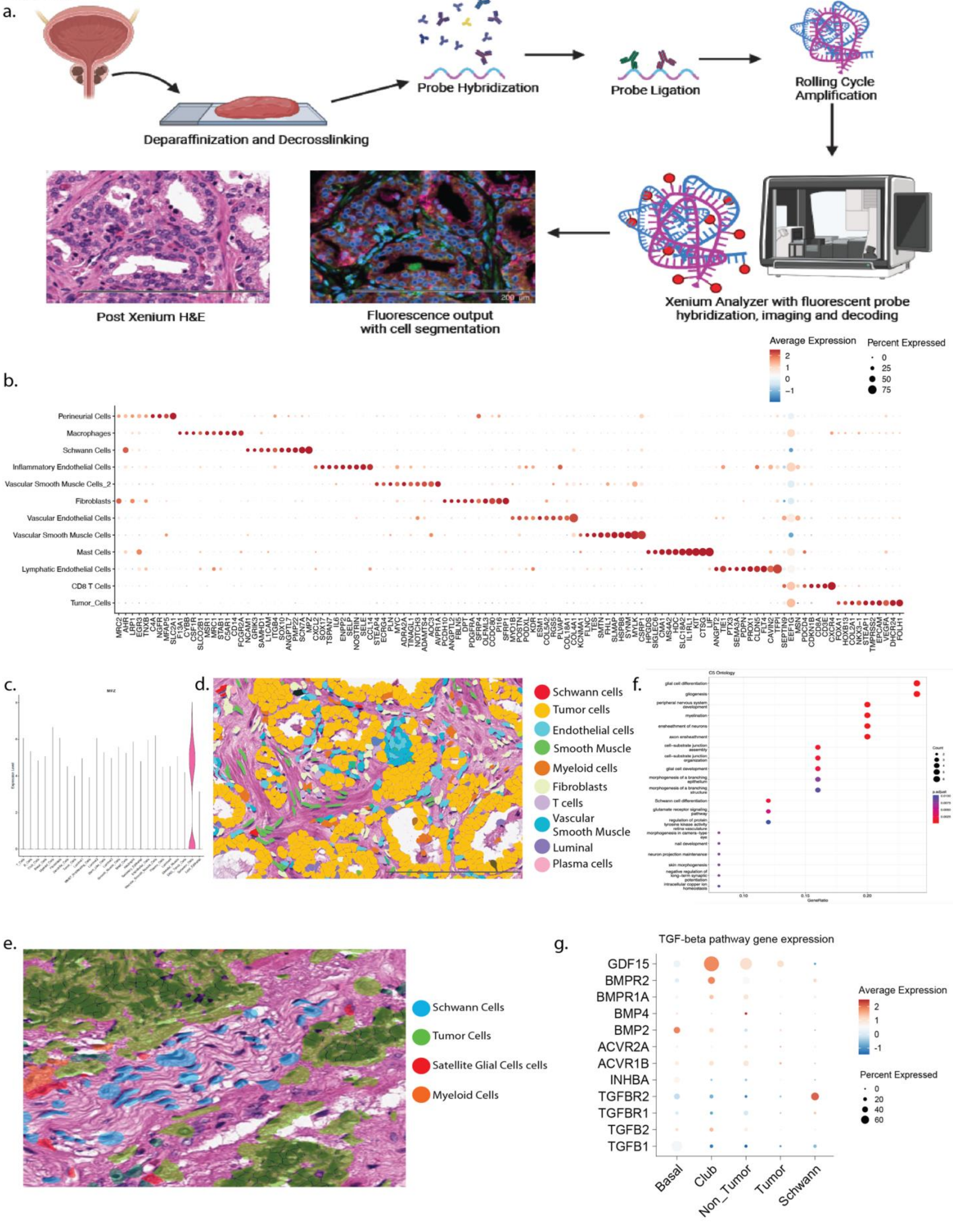

**Supp Figure 8. Characterization of Schwann cells in Xenium spatial data.**

**a.** Overview of the Xenium 5K platform workflow. Tissue sections were formalin-fixed, paraffin-embedded (FFPE), followed by deparaffinization, probe hybridization, ligation, and rolling circle amplification. H&E and fluorescence images illustrate spatial transcriptomic readout and cell segmentation. **b.** Dot plot of selected marker genes for all major annotated cell types in the EUR sample. **c.** Violin plot and cell count bar plot showing Schwann cell abundance and total cell count. **d.** Annotated H&E image from the EUR sample showing Schwann cell distribution relative to tumor glands, stromal, and vascular compartments. **e.** H&E image illustrating the location of Schwann cells (blue) with satellite glial-like cells (red) and myeloid cells (green) around perineural regions. **f.** GO biological process enrichment for Schwann cell-specific DEGs. **g.** Dot plot showing TGFbeta signaling markers in epithelial cells and Schwann cells in the Xenium data.

Supp Figure 9

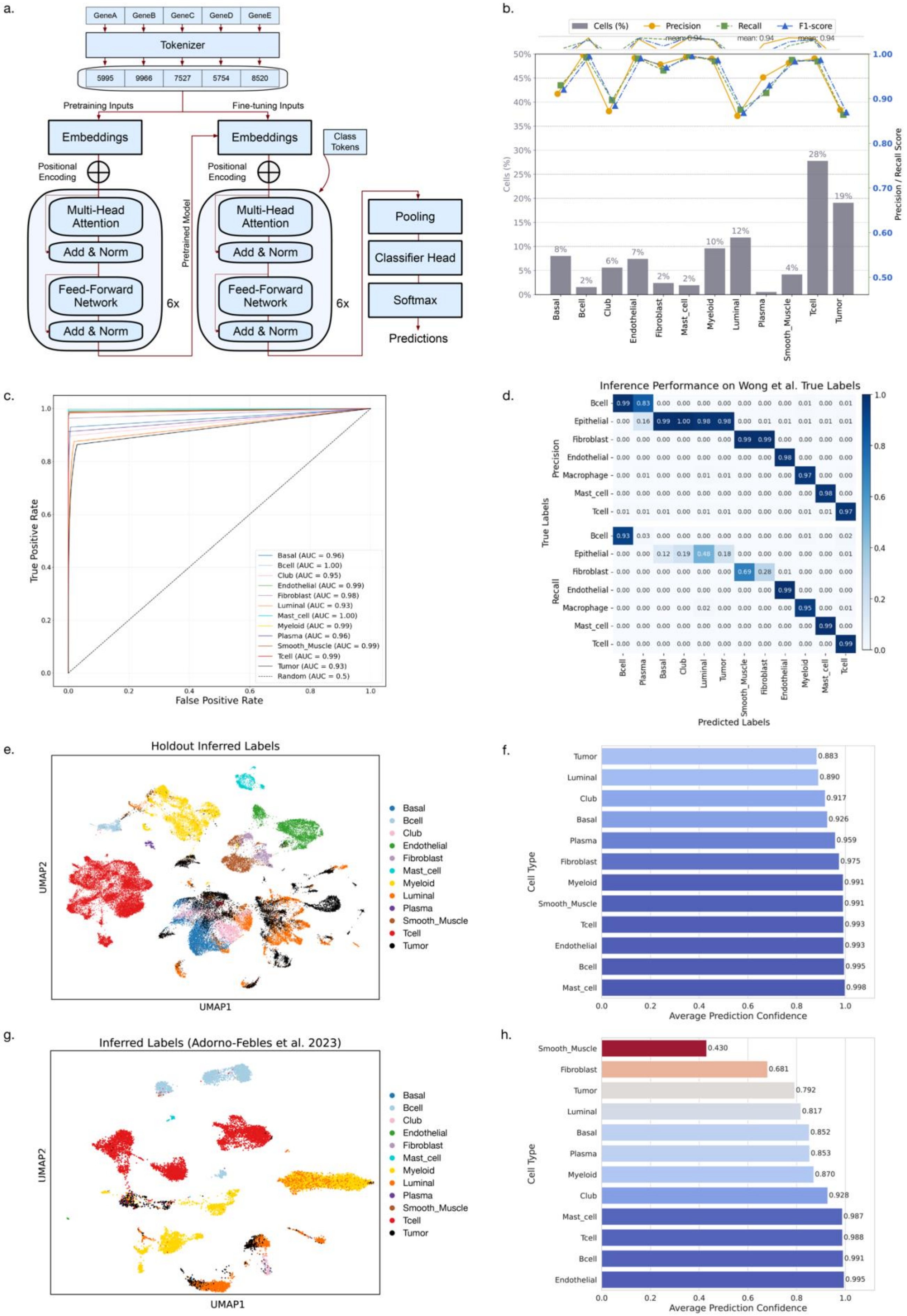

**Supp Figure 9. Evaluation of transformer model inference performance.**

**a.** Flowchart of the deep learning transformer pipeline from data preprocessing to prediction. **b.** Per-class fine-tune performance: precision, recall, and F1-score are high for most cell types (>0.90), with best performance on immune populations (B-cells, mast cells, myeloid cells, plasma cells, T-cells). Misclassifications are most frequent for non-tumor luminal and tumor cells. Orange, green, and blue lines indicate precision, recall, and F1, respectively; gray bars indicate class size. **c.** Receiver operating characteristic (ROC) curves for all predicted Atlas cell types with area under the curve (AUC) > 0.90 across classes. **d.** Model performance on the external Wong et al. scRNA-seq inference dataset, summarized by confusion matrices of precision (top left) and recall (bottom left). **e.** UMAP of predicted cell types in the Atlas hold-out dataset showing well separated clusters corresponding to distinct cell types. **f.** Mean confidence per predicted cell type in the Atlas hold-out dataset. Immune populations are predicted with the highest confidence (>0.90), whereas non-tumor luminal and tumor cells show lower averages (mean confidence 0.89 and 0.88, respectively). **g.** UMAP of predicted cell types for the Adorno-Febles et al. inference dataset, demonstrating well separated clusters and cross dataset generalizability. **h.** Mean confidence per predicted cell type on the Adorno-Febles et al. dataset. Confidence remains high for most lineages, with moderate tumor and non-tumor luminal predictions, and lower fibroblast and smooth muscle confidence scores.
